# Calcium transfer from the ER to other organelles for optimal signaling in *Toxoplasma gondii*

**DOI:** 10.1101/2024.08.15.608087

**Authors:** Zhu-Hong Li, Beejan Asady, Le Chang, Miryam Andrea Hortua Triana, Catherine Li, Isabelle Coppens, Silvia N.J. Moreno

**Author notes:** To whom correspondence should be addressed: Silvia N. J. Moreno, Department of Cellular Biology and Center for Tropical and Emerging Global Disease, 350A Paul D. Coverdell Center, University of Georgia, Athens, GA 30602.

## Abstract

Ca^2+^ signaling in cells begins with the opening of Ca^2+^ channels in either the plasma membrane (PM) or endoplasmic reticulum (ER), leading to a sharp increase of the physiologically low (<100 nM) cytosolic Ca^2+^ level. The temporal and spatial regulation of Ca²⁺ is crucial for the precise activation of key biological processes. In the apicomplexan parasite *Toxoplasma gondii*, which infects approximately one-third of the global population, Ca²⁺ signaling governs essential aspects of the parasite’s infection cycle. *T. gondii* relies on Ca²⁺ signals to regulate pathogenic traits, with several Ca²⁺-signaling components playing critical roles. Ca^2+^ entry from the extracellular environment has been demonstrated in *T. gondii* for both, extracellular parasites, exposed to high Ca^2+^, and intracellular parasites, which acquire Ca²⁺ from host cells during host Ca²⁺ signaling events. Active egress, an essential step of the parasite’s infection cycle, is preceded by a large increase in cytosolic Ca^2+^, most likely initiated by release from intracellular stores. However, extracellular Ca^2+^ is also necessary to reach a cytosolic Ca^2+^ threshold required for timely egress. In this study, we investigated the mechanism of Ca²⁺ intracellular store replenishment and identified a central role for the SERCA-Ca^2+^-ATPase in maintaining Ca²⁺ homeostasis not only within the ER but also in other organelles. We demonstrate mitochondrial Ca^2+^ uptake, which occurs by transfer of Ca^2+^ from the ER, likely through membrane contact sites. Our findings suggest that the *T. gondii* ER plays a key role in sequestering and redistributing Ca²⁺ to intracellular organelles following Ca²⁺ influx at the PM.

**HIGHLIGHTS:** The *T. gondii* ER takes up Ca^2+^ that enters the cytosol from the extracellular milieu.

Filling of acidic stores in *T. gondii* depends on ER Ca²⁺ content.

The mitochondrion of *T. gondii* has no direct access to extracellular Ca²⁺ but can take it up via transfer from the ER and/or acidic stores.

The absence of SERCA activity results in reduced Ca²⁺ levels in the ER as well as in other organelles

## INTRODUCTION

*Toxoplasma gondii* is an intracellular parasite from the Apicomplexan Phylum that infects approximately one third of the world population (Weiss and Dubey, 2009). During the initial infection, *T. gondii* undergoes multiple rounds of a lytic cycle, which consists of host cell invasion, replication within a parasitophorous vacuole (PV), exit from the host cell causing its lysis followed by reinvasion of new host cells (Black and Boothroyd, 2000; Blader *et al*., 2015). Cytosolic Ca^2+^ ([Ca^2+^]c) fluctuations precede the activation of several key steps of the *T. gondii* lytic cycle like motility, attachment, invasion, and egress (Lourido and Moreno, 2015; Hortua Triana *et al*., 2018). Egress from the host cell is an essential step for the infection cycle of *T. gondii* (Bisio and Soldati-Favre, 2019) and it was shown that it is preceded by a cytosolic Ca^2+^ increase (Endo *et al*., 1982; Borges-Pereira *et al*., 2015). Extracellular Ca^2+^ entry was demonstrated in extracellular (Pace *et al*., 2014; Hortua Triana *et al*., 2024) and intracellular replicating tachyzoites (Vella *et al*., 2021). This activity was highly regulated and work from our lab revealed that a TRP-like activity was involved (Marquez-Nogueras *et al*., 2021).

Ca^2+^ signaling is part of the signaling pathways that regulate a large number of cellular functions (Clapham, 2007). All cells express a variety of channels, transporters, and Ca^2+^ pumps, located at the plasma membrane (PM) and/or intracellular organelles (endoplasmic reticulum (ER), acidic stores, and mitochondria) that regulate/control the concentration of cytosolic Ca^2+^. However, an elevated cytosolic Ca²⁺ concentration sustained for prolonged periods is toxic to cells and may result in their death (Bootman and Bultynck, 2020).

In *T. gondii*, both Ca^2+^ entry through the plasma membrane and release from intracellular stores like the ER may initiate a cascade of signaling events important for the stimulation of the biological steps of the parasite lytic cycle (Lourido and Moreno, 2015; Hortua Triana *et al*., 2018). Ca^2+^ oscillations were observed in motile parasites loaded with fluorescent Ca^2+^ indicators, (Lovett and Sibley, 2003) as well as expressing Genetically Encoded Calcium Indicators (GECIs) (Borges-Pereira *et al*., 2015). The significance of Ca^2+^ signals during all stages of the lytic cycle has been demonstrated, but little is known about the mechanism by which intracellular stores contribute to cytosolic Ca^2+^ signals and downstream regulation.

The ER, an exclusive organellar feature of eukaryotic cells, is the main store for Ca^2+^ in most eukaryotic cells. It has been proposed that the ER is functionally heterogeneous with Ca^2+^ binding proteins and Ca^2+^ pumps and channels having a nonuniform distribution resulting in the presence of distinct subdomains within the organelle (Papp *et al*., 2003). The ER in mammalian cells facilitates Ca^2+^ tunneling through its lumen as a mechanism of delivering Ca^2+^ to targeted sites without activating inappropriate processes in the cell cytosol (Petersen *et al*., 2017). In addition, the ER is ubiquitously distributed and is in close contact with all cellular organelles and the plasma membrane (PM) (Spang, 2018). Over the past decade a new paradigm has emerged that seeks to decipher how subcellular organelles communicate with each other in order to coordinate activities and efficiently distribute ions and lipids within the cell. Numerous observations have highlighted the presence of tight, stable and yet non-fusogenic associations between organellar membranes which have since become known as membrane contact sites (MCSs) (Phillips and Voeltz, 2016).

The secretory pathway of *T. gondii* is organized in a highly polarized manner with the ER being an extension of the nuclear envelope (Hager *et al*., 1999; Tomavo *et al*., 2013). The ER at the apical surface of the nuclear envelope continues with the Golgi stacks and then the secretory organelles, micronemes and rhoptries, which are unique to the apicomplexan phylum (Hager *et al*., 1999). These organelles perform important functions relevant to a successful lytic cycle like host cell attachment, invasion, and establishment of the parasitophorous vacuole (PV). Cytoplasmic Ca^2+^ increases, due to release from the ER, have been reported to initiate responses like microneme secretion (Carruthers and Sibley, 1999; Nagamune *et al*., 2007b), conoid extrusion (Del Carmen *et al*., 2009), invasion (Vieira and Moreno, 2000; Lovett and Sibley, 2003) and egress (Arrizabalaga and Boothroyd, 2004). These responses require precise spatiotemporal control of diverse targets and suggests the presence of distinct systems to deliver Ca^2+^ to specific locations rather than allowing global increases, which would activate unnecessary and potentially detrimental signaling events (Huet and Moreno, 2023).

In order to concentrate Ca^2+^ ions, the ER utilizes a SERCA-Ca^2+^-ATPase, a transmembrane P-type ATPase, that couples ATP hydrolysis to the transport of ions across biological membranes and against a concentration gradient. SERCA pumps can be inhibited by various inhibitors including the very potent and highly specific thapsigargin (TG) (Thastrup *et al*., 1990; Sagara and Inesi, 1991).

*T. gondii* expresses a SERCA Ca²⁺-ATPase (TgSERCA), which possesses conserved SERCA domains, Ca²⁺-binding sites, and residues required for ATP hydrolysis (Nagamune *et al*., 2007a). The function of TgSERCA was determined by rescue experiments of yeast cells defective in Ca^2+^-ATPases and by its specific inhibition by TG (Nagamune *et al*., 2007a). TgSERCA was mainly localized to the ER of *T. gondii* but also showed a distinct distribution in extracellular parasites, where the protein was partially found in ER vesicles in the apical region near micronemes (Nagamune *et al*., 2007a). This distribution pattern was different to the one obtained with the transient transfection of GFP-HDEL (an ER marker), which was retained near the nuclear envelope, suggesting an uneven distribution of ER markers in extracellular parasites. The authors suggested that this distribution to the apical end may be important for rapid release and effective recovery of cytosolic Ca^2+^, events that likely govern both motility and microneme secretion (Nagamune *et al*., 2007a; Nagamune *et al*., 2007b).

In this work, we investigate the role of the ER in intracellular calcium handling in *Toxoplasma gondii*. Specifically, we explore how the ER contributes to calcium uptake following extracellular influx and how it facilitates calcium redistribution to other organelles. Using genetic and pharmacological tools, we examine the activity of the SERCA Ca²⁺-ATPase and its role in coordinating calcium dynamics. Our findings support a model in which the ER acts as a central hub for calcium buffering and trafficking, driven by the high calcium affinity of TgSERCA.

## RESULTS

### The ER sequesters and redistributes Ca^2+^ to other organelles following PM influx

We initially designed experiments to track the destination of Ca^2+^ taken up by extracellular tachyzoites from the extracellular milieu into their cytosol. Tachyzoites were loaded with the ratiometric Ca^2+^ indicator Fura-2 (**Fig. 1A**) and incubated in Ringer buffer containing 100 µM EGTA to chelate extracellular Ca^2+^ and prevent further uptake. Under these conditions, we investigated the release of intracellular Ca^2+^ pools by various pharmacological agents.

**Figure 1:**
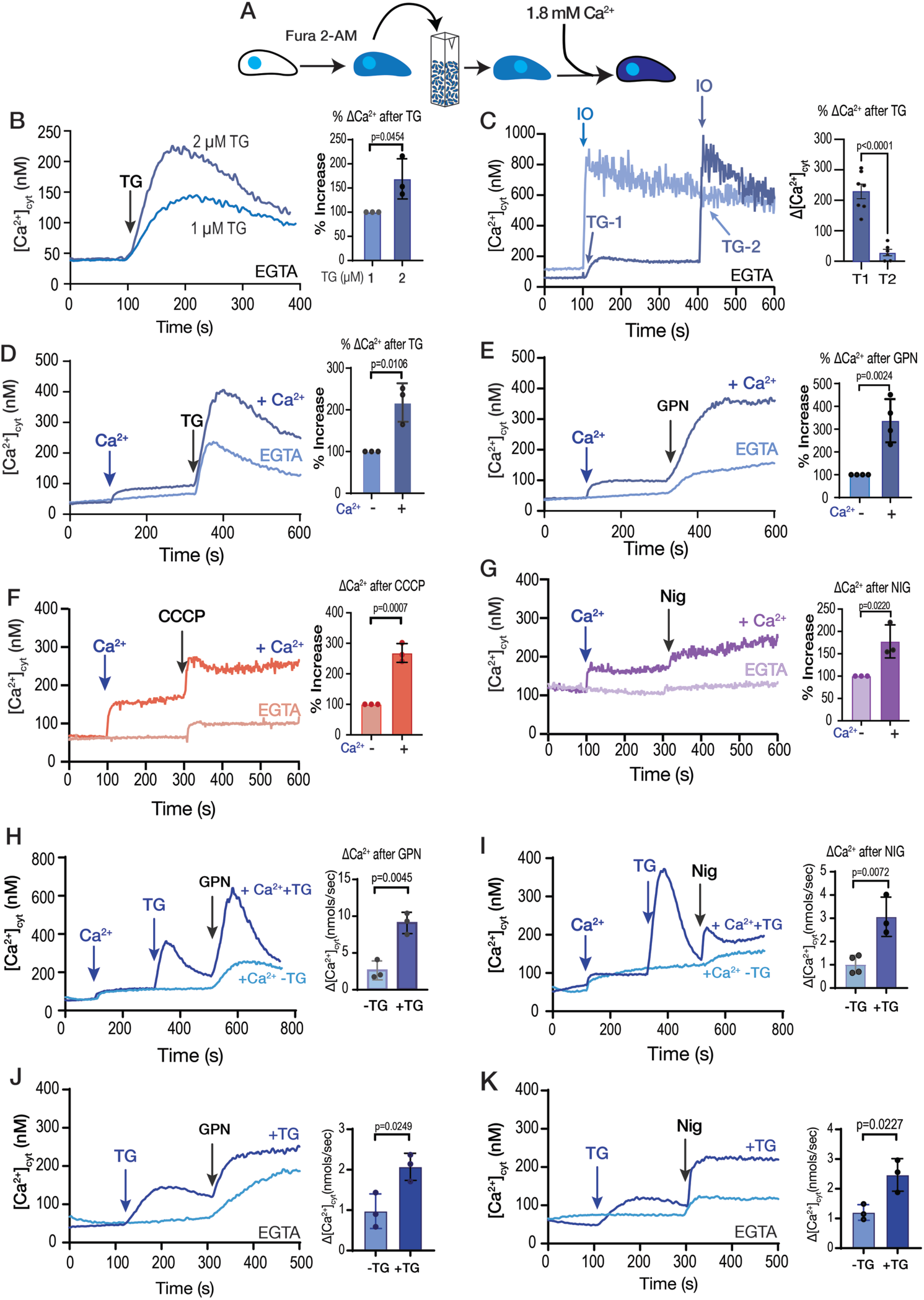
The role of extracellular calcium in the filling of intracellular stores. **A**, scheme depicting the Fura-2-AM loading and the experimental set-up. **B**, *T. gondii* tachyzoites loaded with Fura-2 were in suspension in Ringer buffer with 100 µM EGTA. Thapsigargin (TG) was added at 100 seconds, at two different concentrations (1 and 2 µM). **C**, Same conditions as in B. 1 µM Thapsigargin (TG) was added at 100 sec for T1 and 400 sec for T2. 1 µM ionomycin (IO) was added at 400 sec for trace 1 (T1) and at 100 sec for trace 2 (T2). Bar graph shows Ca^2+^ increase after adding TG before (T1) and after IO (T2). **D**, Same conditions as in B. 1.8 mM CaCl2 was added at 100 sec, followed by 2 µM TG at 300 sec (dark blue trace). The light blue trace shows the same experiment without the addition of CaCl2. **E**, similar to D but using 40 µM GPN instead of TG. **F,** same experimental set-up to the one shown in D but using the mitochondrial uncoupler CCCP. **G,** same experimental set-up to the one shown in D but adding the potassium ionophore nigericin (Nig), 10 µM. The quantification for D, E, F, and G shows the % increase of cytoplasmic calcium compared with the same condition without calcium. **H,** 1.8 mM CaCl2 was added at 100 sec, 1 µM TG was added at 300 sec followed by 40 µM GPN at 500 sec. The light blue trace shows the same experiment without the addition of TG. The quantification shows the Ca^2+^ increase after adding GPN ± previous addition of TG**. I,** Identical experiment to H but instead using 10 µM Nig at 500 sec. The quantification shows the Ca^2+^ increase after adding Nig ± previous addition of TG. **J,** 1 µM TG was added at 100 sec followed by 40 µM GPN at 300 sec. The light blue trace shows the same experiment without the addition of TG. The buffer contains 100 µM EGTA. Quantification shows Ca^2+^ increase after adding GPN ± previous addition of TG **K,** Similar conditions to J but with10 µM Nig at 300 sec instead. Data are presented as mean ± SD for all comparisons. *p* value: unpaired two tailed t test performed in all comparisons.

When thapsigargin (TG), an inhibitor of SERCA, was added, it blocked the re-uptake of Ca²⁺ into the ER and unmasked the passive leak of Ca²⁺ from the ER into the cytosol, resulting in a cytosolic Ca²⁺ increase of approximately 160 nM **(Fig. 1B-C)**. We compared this response to the effect of ionomycin (IO), a Ca^2+^/H^+^ ionophore which acts on neutral intracellular Ca^2+^ stores, inducing depletion of Ca^2+^, mainly from the ER (Smith *et al*., 1989). Exposure of *T. gondii* to 1 µM IO caused a cytosolic increase of approximately 700-1100 nM Ca^2+^ (**Fig. 1C**). We next tested another SERCA inhibitor, cyclopiazonic acid (CPA), which is structurally unrelated to TG and with a different mode of action (Inesi and Sagara, 1994). CPA induced a smaller increase in cytosolic Ca²⁺ compared to TG, possibly indicating less efficacy toward TgSERCA **(Fig. S1A-B).** Exposure to TG before CPA abolished the effect of CPA, whereas exposure to CPA did not prevent the effect of TG. **(Fig. S1B-C)**. Although TG does a better job at depleting the ER of Ca^2+^ in intact parasites, the resulting increase of cytosolic Ca^2+^ after adding TG is modest compared to the response of IO **(Fig. 1C)**. This result could reflect the slow kinetics of Ca²⁺ leak from the ER, allowing other buffering and transport mechanisms to mitigate the phenomenon. Alternatively, it may indicate the duration after TG treatment allowing time to complete store depletion. As shown in **Fig. 1B-C**, residual Ca²⁺ remains in the stores after TG treatment, and the TG-induced phenomenon does not return to baseline, suggesting that the leak remains active.

Next, we tested the effect of Ca^2+^ entry in the filling of intracellular stores by measuring the cytosolic Ca^2+^ increases in response to inhibitors after pre-exposing the parasite suspension to extracellular Ca^2+^ (**Fig. 1D-G**, *darker traces*). Addition of 1.8 mM Ca^2+^ caused a cytosolic increase due to influx through the plasma membrane (**Fig. 1D**, *dark blue trace*) (Pace *et al*., 2014). A substantial portion of the entering Ca²⁺ appeared to be rapidly sequestered by the ER, as evidenced by the significantly greater TG-induced response in parasites previously exposed to extracellular Ca²⁺ (compare light and dark blue traces) (**Fig. 1D**). We next tested the lysosomotropic agent glycyl-L-phenylalanine-naphthylamide (GPN), which primarily mobilizes Ca^2+^ from acidic organelles (Haller *et al*., 1996; Lloyd-Evans *et al*., 2008; Miranda *et al*., 2010; Yuan *et al*., 2021) and observed a similar pattern (**Fig. 1E**). The increase in cytosolic Ca²⁺ following GPN addition was markedly greater in parasites previously exposed to extracellular Ca²⁺ than in those that had not been exposed (**Fig. 1E**, *compare light and dark blue traces*). We previously proposed that GPN may act on the lysosome-like Plant-Like Vacuolar Compartment (PLVAC), a dynamic acidic organelle involved in calcium storage, and the processing of secretory proteins (Miranda *et al*., 2010);(Stasic *et al*., 2022). The increase in cytosolic Ca²⁺ in response to the addition of nigericin (which acts on acidic stores) or CCCP (which likely targets mitochondria) was also greater in cells previously loaded with extracellular Ca^2+^ (**Fig. 1F-G**, *dark purple and orange traces, respectively*). These data indicate that the ER, mitochondrion, PLVAC, and other acidic stores release more calcium into the cytosol of *T. gondii* tachyzoites following exposure to extracellular calcium, which stimulates its influx through the plasma membrane. The ER displayed high capacity to access a large portion of extracellular Ca^2+^, with TG producing close to ∼300-400 nM of Ca^2+^ increase after pre-exposure to Ca^2+^ and only ∼150-200 nM Ca^2+^ without Ca^2+^ pre-exposure **(Fig. 1D)**.

We next aimed to understand how other compartments are replenished with Ca²⁺, given that the ER appears to be particularly effective at taking up Ca²⁺ from the cytosol. We designed an experiment where parasites were first loaded with Ca²⁺, followed by inhibition of SERCA using TG. This inhibition prevents ER Ca²⁺ uptake, allowing Ca²⁺ to accumulate on the cytosolic side of the ER membrane. Under these conditions, we added agonists such as GPN or nigericin following the addition of TG. As shown in **Fig. 1H**, Ca²⁺ was first added to load intracellular stores, followed by TG to induce ER Ca²⁺ leakage, and then GPN to trigger Ca²⁺ release from acidic stores. Comparison of GPN-induced cytosolic Ca²⁺ signals with and without TG pre-treatment revealed a significantly greater response in the TG condition. A similar enhancement was observed for the nigericin-induced response following TG treatment (**Fig. 1I**).

These results support our hypothesis that extracellular Ca²⁺ is primarily taken up by the ER and subsequently redistributed to other organelles. Importantly, Ca²⁺ was added prior to TG to allow store loading, and TG treatment then permitted ER Ca²⁺ leakage, facilitating Ca²⁺ transfer to other compartments.

However, we considered the possibility that the enhanced responses to GPN or nigericin could be due to increased PM Ca²⁺ influx triggered by elevated cytosolic Ca²⁺. To test this, we repeated the experiments in the absence of extracellular Ca²⁺ (**Fig. 1J-K**). Notably, prior addition of TG resulted in an enhanced cytosolic Ca²⁺ response to both GPN and nigericin. These results further support the notion that Ca²⁺ can be transferred from the ER to other intracellular stores independently of extracellular Ca²⁺ influx.

We also performed an additional experiment in which SERCA was inhibited with TG prior to Ca²⁺ addition. We then quantified the subsequent GPN response in conditions with and without TG preincubation and observed a significant increase in the TG-treated group (**Fig. S2A**). This result suggests that, under non-physiological conditions where SERCA is blocked, the PLVAC may take up Ca²⁺ directly from the cytosol. However, this is unlikely to occur under normal conditions, as functional SERCA likely has a higher affinity for Ca²⁺ and would sequester it limiting its availability to other compartments.

In summary, pre-exposure of *T. gondii* to physiological levels of extracellular Ca²⁺ markedly enhanced the capacity of the ER, mitochondria, and acidic stores to release calcium, with the ER and GPN-sensitive stores exhibiting the most pronounced responses.

### Ca^2+^ uptake by the SERCA-Ca^2+^ ATPase in permeabilized tachyzoites

The previous results highlight the central role of the ER in taking up Ca²⁺ from the cytoplasm following an influx from the extracellular milieu. We propose that this ER uptake, essential for maintaining Ca²⁺ store levels, is driven by the high Ca²⁺ affinity of TgSERCA. As a key mechanism, SERCA enables the ER to sustain its Ca²⁺ concentration despite the constitutive and passive leakage of Ca²⁺ from the ER into the cytosol (Camello *et al*., 2002).

To characterize the activity of TgSERCA *in situ*, we adapted a protocol to directly measure Ca²⁺ uptake by the stores in which TgSERCA localizes (ER and Golgi apparatus) (Calixto *et al*., 2025). This approach, which has been widely used in mammalian cells to assess Ca²⁺ release from the ER, employs the low-affinity Ca²⁺ indicator Mag-Fluo-4 (Kd ∼22 µM) (Rossi and Taylor, 2020). The cytosolic concentration of Ca^2+^ in *T. gondii* is approximately 70 nM (Moreno and Zhong, 1996), which is well below the detection threshold of Mag-Fluo-4. To facilitate loading into organelles, we incubated parasites for an extended period with higher concentrations of Mag-Fluo-4-AM, promoting its compartmentalization into intracellular stores. Following incubation, parasites were washed and treated with a low concentration of digitonin, which selectively permeabilizes the plasma membrane while preserving the integrity of organellar membranes. Under these conditions, the parasites retained the Ca²⁺ indicator within their organelles (**Fig. 2A**). We next assessed the capacity of these permeabilized parasites to take up Ca^2+^. Since the activity of SERCA depends on MgATP (**Fig. 2B**), we added this substrate in the presence of defined Ca^2+^ concentrations calculated using the MaxChelator program (Bers *et al*., 1994). Under these conditions (free calcium ranging from 55-880 nM and MgATP at concentrations of 25-500 μM), we observed consistent and reproducible Ca^2+^ uptake, as shown in **Fig. 2B-C**. We selected 220 nM Ca^2+^ for our study because this concentration approximates physiological cytosolic fluctuations and supports detectable Ca^2+^ uptake. Additionally, this concentration of Ca^2+^ has been used in previous studies of mammalian SERCA (Rossi and Taylor, 2020). Validation that this activity is mediated by TgSERCA is demonstrated by the addition of TG, which inhibits SERCA allowing Ca²⁺ leakage from the organelle (**Fig. 2D**). Although the Ca²⁺ released after adding TG appears modest, consistent with the slow leak characteristics of ER calcium, the high Kd (22 µM) of the indicator implies that even small decreases in fluorescence signal, represent significant Ca²⁺ efflux. In contrast, IO at 1 µM caused a more pronounced Ca²⁺ release, lowering the Ca²⁺ concentration below the baseline level. This effect is likely due to IO targeting multiple intracellular compartments in addition to the ER, as well as the fundamental difference in mechanisms: IO acts as an ionophore, directly facilitating Ca²⁺ efflux across membranes, whereas TG inhibits SERCA, resulting in Ca²⁺ release through the ER’s natural leak pathway (**Fig. 2E**).

**Figure 2:**
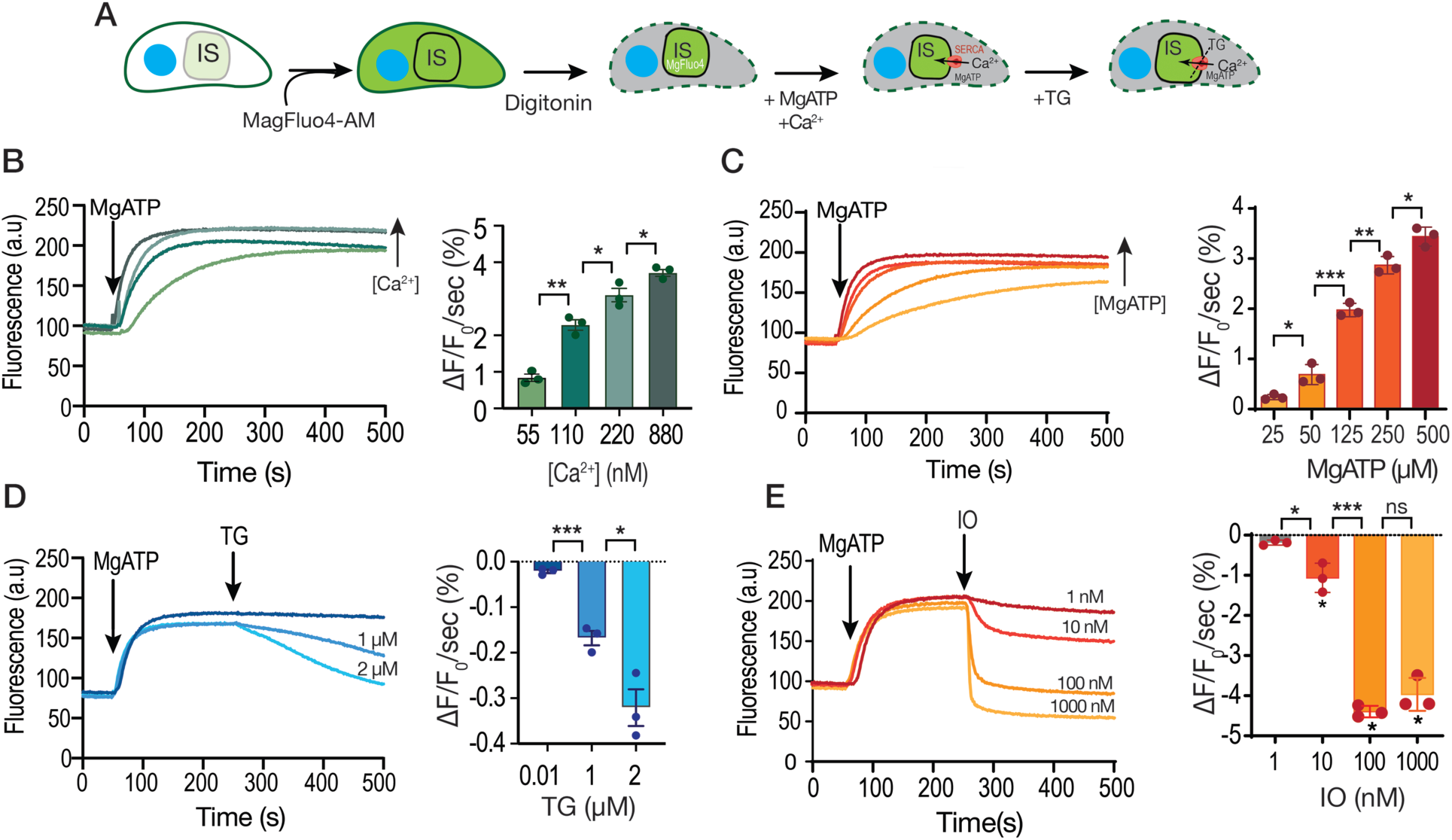
Ca^2+^ uptake by intracellular stores. A, Scheme showing the loading with Mag-Fluo-4 AM followed by permeabilization with digitonin of a *T. gondii* tachyzoite (RH parental strain) suspension (IS, intracellular store). **B,** Fluorescence measurements (see Materials and Methods for specifics) of the suspension of parasites loaded with Mag-Fluo-4. MgATP (500 µM), the SERCA substrate was added at 50 seconds. The bar graph shows the quantification of the slope of the increase in fluorescence after adding MgATP. The concentration of free calcium was varied, and it is indicated. The calculation of free calcium was done using MaxChelator. **C,** similar experimental set-up to the one shown in B with 220 nM free Ca^2+^, with varied concentrations of MgATP as indicated in the bar graph, which shows the quantification of the slope of fluorescence increase after adding MgATP. **D,** experiment was done with 500 µM MgATP and 220 nM free Ca^2+^. Thapsigargin (TG) was added to inhibit SERCA causing calcium to be released from the store. The concentrations used are indicated. The bar graph shows the negative slope after the addition of TG. **E,** similar to D, but adding various concentrations of ionomycin (IO). The concentrations used are indicated and the slopes were measured after the addition of IO. Data are presented as mean ± SD for B-D. *p* value: unpaired two tailed t test performed in all comparisons. ns, not significant, p > 0.05. *, p ≤ 0.05. **, p ≤ 0.01. ***, p ≤ 0.001. ****, p ≤ 0.0001

We used the Mag-Fluo-4 assay to directly compare the inhibitory effects of CPA and TG (**Fig. S3)**. Under the conditions of the Mag-Fluo-4 assay, using digitonin-permeabilized parasites, both inhibitors produced comparable levels of Ca^2+^ efflux suggesting that at the concentrations used both inhibited SERCA and the efflux rate corresponds to the intrinsic ER leak mechanism (**Fig. S3A–C**). This finding suggests that CPA may be less effective at inhibiting SERCA in intact parasites, possibly due to its reversibility and partial dissociation over time, allowing residual Ca²⁺ reuptake into the ER and resulting in a smaller cytosolic Ca²⁺ increase compared to TG.

In summary, these results demonstrate that the activity of TgSERCA in *T. gondii* tachyzoites can be measured *in situ* using permeabilized parasites loaded with the low-affinity Ca²⁺ indicator Mag-Fluo-4. This activity is MgATP-dependent and both TG and CPA can inhibit TgSERCA activity, leading to leakage of the accumulated Ca²⁺. The larger effect of IO compared to TG is likely due to differences in their mechanisms of action.

### TgSERCA and the *T. gondii* lytic cycle

To investigate the role of TgSERCA (TGGT1_230420) in the biology of *T. gondii* we generated conditional knockout parasites (*i!ιTgSERCA*), based on the gene’s predicted essentiality (fitness score -5.44) (Sidik *et al*., 2016). A tetracycline regulatable element was inserted at the 5’ end of the *TgSERCA* gene to control its expression with anhydrotetracycline (ATc) (Sheiner *et al*., 2011). In addition, we endogenously tagged TgSERCA with a C-terminal 3xHA epitope and generated clonal lines of both *iΔTgSERCA* and *iΔTgSERCA-3HA* (**Fig. 3A**).

**Figure 3:**
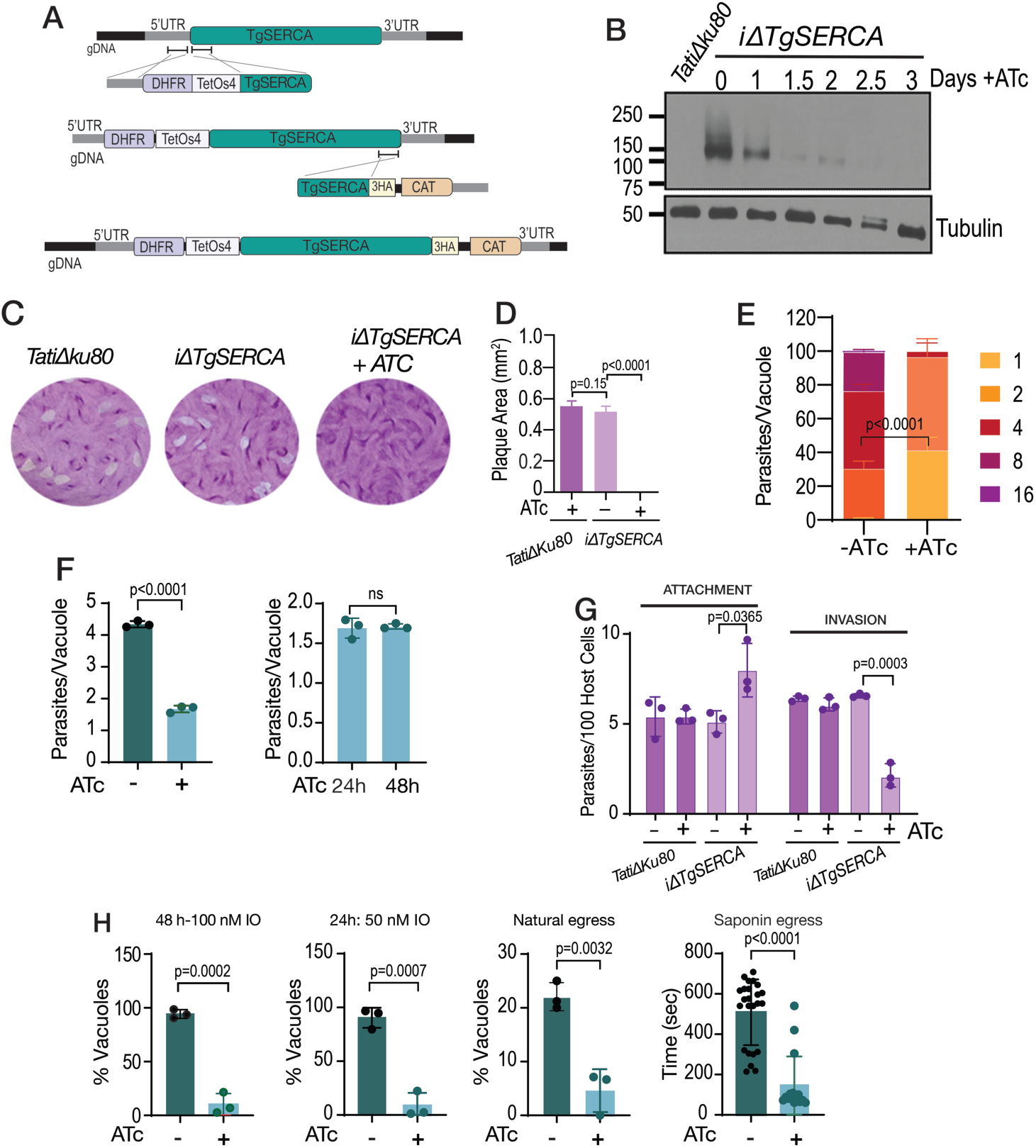
The sarcoplasmic-endoplasmic reticulum calcium ATPase (SERCA) is essential for the *T. gondii* lytic cycle. **A**, Scheme showing the strategy used for generating conditional knock outs of TgSERCA by promoter insertion and regulation by 0.5 µg/ml Anhydrotetracyclin (ATc). The resulting mutants were named *iΔTgSERCA* or *iΔTgSERCA-3HA* (C-terminally HA-tagged). DHFR, dihydrofolate reductase gene (pyrimethamine selection); CAT, chloramphenicol acetyltransferase gene (chloramphenicol selection). **B**, Western blots of *iΔTgSERCA-3HA* parasites grown ± ATc. TgSERCA expression was detected using an anti-HA antibody, showing reduced levels with ATc treatment. **C**, plaque assays comparing the growth of *iΔTgSERCA* tachyzoites (150 parasites/well) cultured ± 0.5 µg/ml ATc for 8 days. Plaques formed by the parental *TatiΔku80* strain are shown for comparison. **D**, quantification of the size of the plaques presented in C. **E**, Replication assay using the *iΔTgSERCA-RFP* mutant. The number of parasites per parasitophorous vacuole (PV) was quantified 24 h post-infection of fibroblast cells and compared between parasites grown ± 0.5 µg/ml ATc. **F**, Average number of parasites per PV counted at 24 h after the initial infection. The graph to the right shows the number of parasites per PV of the *iΔTgSERCA* (+ATc) for 24 or 48 hours after the initial infection. **G**, Invasion assay of the *iΔTgSERCA* mutant following 24 h of ATc treatment, performed using the red-green assay described in Materials and Methods. **H,** Egress assays with fibroblast monolayers infected with *iΔTgSERCA-RFP* parasites for 24 or 48 h. Egress was triggered with ionomycin (IO; 100 nM or 50 nM) or saponin (0.01%). Natural egress was monitored following treatment with 1 μM compound 1 as described in the Methods section. % Vacuoles: 100 X Vacuoles egressed/total vacuoles. Data (D, E, F, G, H) are presented as mean from at least three biological replicates ± SD. Statistical significance was assessed using an unpaired two-tailed t-test.

With the aim to detect the protein, we generated a guinea pig polyclonal antibody against the phosphorylation (P) and nucleotide-binding (N) domains of TgSERCA, which was affinity-purified and validated by Western blotting (**Fig. 3B**) and IFAs (**Fig. S4**). Colocalization of TgSERCA with HA epitope was confirmed by IFA. Although the signals from the anti-HA and anti-TgSERCA did not completely overlap, both were lost in the *iΔTgSERCA* mutant grown in the presence of ATc (**Fig. S4,** *+ATc*). The partial colocalization may reflect differences in antibody accessibility or that the two antibodies recognize distinct regions of the protein. Both Western blots and IFAs confirmed that TgSERCA expression is tightly regulated by ATc and becomes undetectable after 2.5 days in culture (**Fig. 3B** **and S4**). Growth of the *iΔTgSERCA* mutant was severely impaired in the presence of ATc, as assessed by plaque assays (**Fig. 3C–D**). In this assay, parasites undergo successive rounds of invasion, replication, and egress, leading to host cell lysis and the formation of plaques on confluent monolayers. Downregulation of TgSERCA expression led to a marked defect in replication, with parasites failing to progress beyond one or two rounds of division (**Fig. 3E–F**). All parasitophorous vacuoles (PVs) in ATc-treated cultures contained four or fewer parasites (**Fig. 3F**, *bar graph on the right*). Host cell invasion was also reduced in the *iΔTgSERCA* mutant (**Fig. 3G**) when cultured with ATc.

Parasite egress was significantly affected by TgSERCA depletion. Ionomycin (IO), which has been known to trigger egress by inducing Ca²⁺ release (Borges-Pereira *et al*., 2015), and natural egress following pre-incubation with 1 μM compound 1 (Donald *et al*., 2002), previously shown to synchronize parasite exit (Vella *et al*., 2021), were both markedly reduced in ATc-treated parasites, underscoring the critical role of ER Ca²⁺ stores in supporting both ionophore-induced and spontaneous egress (**Fig. 3H**, *IO and natural egress*). Interestingly, however, egress induced by saponin in the presence of extracellular Ca²⁺ was accelerated in ATc-treated parasites (**Fig. 3H**, *saponin egress*). This enhancement may result from a more rapid rise in cytosolic Ca²⁺, reaching the egress threshold more quickly due to impaired SERCA activity combined with ongoing Ca²⁺ leak from the ER (Vella *et al*., 2021). The saponin concentration used selectively permeabilizes the host cell membrane, allowing extracellular Ca²⁺ to enter the parasite cytosol without compromising the integrity of the parasite plasma membrane. This is consistent with previous observations showing that tachyzoites remain motile and exhibit Ca²⁺ oscillations under similar conditions (Borges-Pereira *et al*., 2015). The resulting rise in cytosolic Ca²⁺ within the parasite stimulates motility and triggers egress. To further examine this phenomenon, we directly compared the timing of egress between untreated and ATc-treated *iΔTgSERCA* parasites under identical saponin exposure conditions (**Fig. 3H**).

In summary, our findings demonstrate that TgSERCA is essential for *T. gondii* replication, invasion, and natural egress. Interestingly, when host cells were selectively permeabilized, parasites with reduced TgSERCA expression displayed accelerated egress, likely due to altered calcium dynamics.

### Ca^2+^ uptake by the SERCA-ATPase is essential for filling acidic Ca^2+^ stores

Further characterization of the *i!ιTgSERCA* mutant showed a diminished cytosolic Ca^2+^ response to TG (**Fig. 4A**), which was also observed when TG was applied after extracellular Ca^2+^ had been added to fill the stores (**Fig. 4B**). This was most likely due to reduced Ca^2+^ accumulation by the ER in the *i!ιTgSERCA* (+ATc) mutant. Note that the change in Ca^2+^ in **Fig. 4A** (*Tati!ιku80* cells) is larger than in **Fig. 1B** (RH strain), which we attribute to differences between the two cell lines (RH vs *Tati!ιku80*).

**Figure 4:**
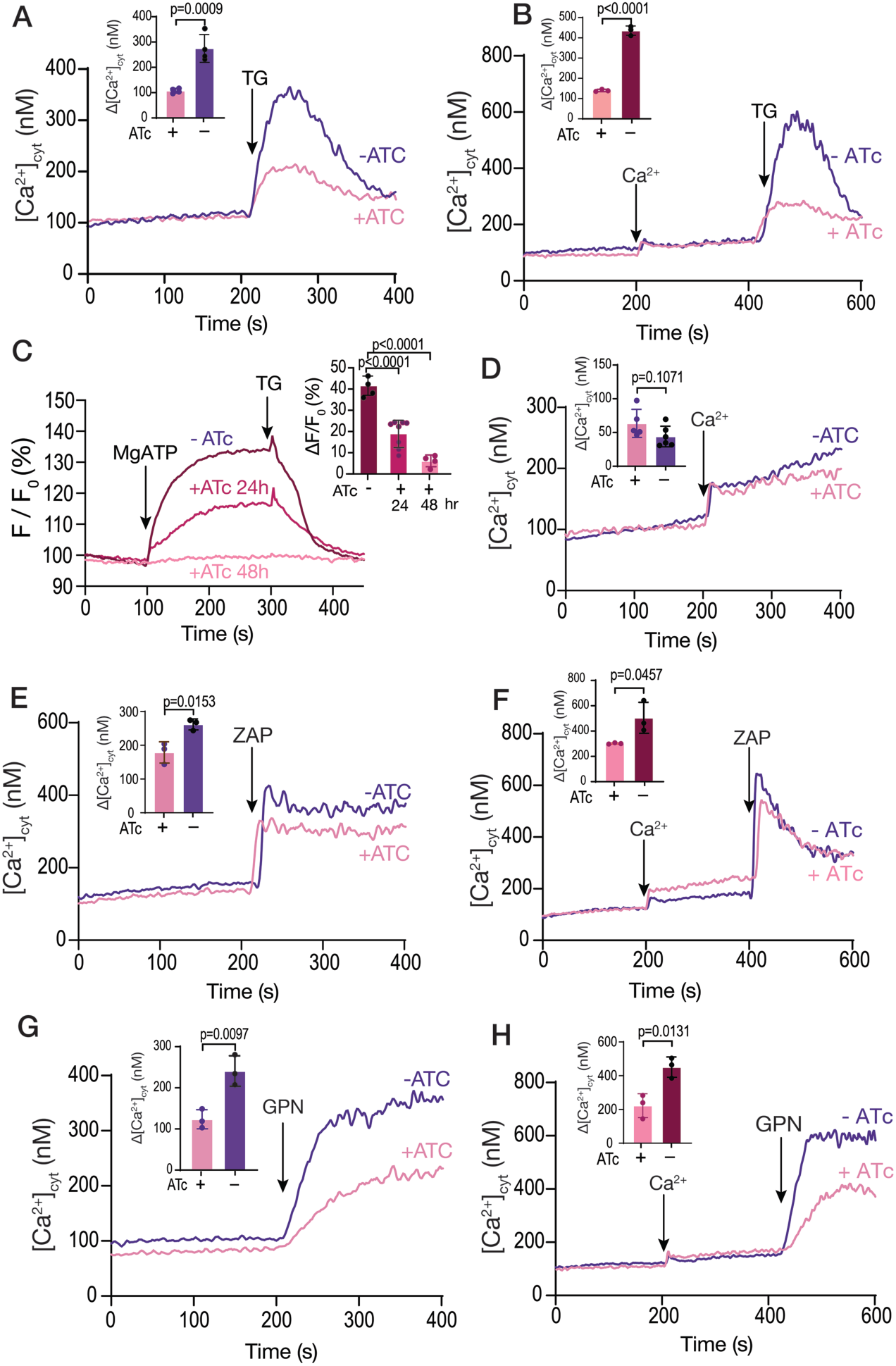
Organellar calcium pools in the *iΔTgSERCA* mutant. A, The *iΔTgSERCA* mutant was grown ± ATc and was loaded with Fura-2 for cytosolic Ca^2+^ measurements. 1 µM TG was added at 200 sec to a suspension of tachyzoites. The purple trace shows the response of the parental cell line grown without ATc and the pink trace shows the response of the same mutant grown with ATc for 24 h. The bar graph shows the analysis of the Δ[Ca^2+^]cyt from three biological experiments. B, same experimental set-up as the one in A but adding 1.8 mM extracellular Ca^2+^ at 200 sec. C, SERCA activity measured in Mag-Fluo-4 loaded *iΔTgSERCA* tachyzoites grown ± ATc. Parasites were collected, loaded with Mag-Fluo-4 AM, and permeabilized with digitonin as described in the Methods section. Free Ca²⁺ in the buffer was set at 220 nM, and MgATP (0.125 mM) was added at 100 s. The purple trace represents the control (no ATc), while the other traces correspond to parasites treated with ATc for 24 or 48 h. TG (1 µM) was added as indicated. The bar graph shows the quantification of the initial slope after adding MgATP. D, Ca²⁺ entry measured in Fura-2–loaded *iΔTgSERCA* parasites grown ± ATc. Extracellular Ca²⁺ (1.8 mM) was added at 200 s. The inset shows ΔF values from three independent experiments, indicating no significant differences. E, Similar conditions to the ones used in A but adding 100 µM Zaprinast. The bar graph shows the quantification of the D[Ca^2+^] from three biological experiments. F, Similar conditions to the ones used in B but adding 1.8 mM extracellular calcium at 200 sec and 100 µM Zaprinast at 400 sec. The bar graph shows the quantification of the Δ[Ca^2+^] from 3 biological experiments. G, Same as A but adding 40 µM GPN. The bar graph shows the analysis of the Δ[Ca^2+^] from three biological replicates. H, Same set-up as in F but adding 1.8 mM Ca^2+^ at 200 sec followed by 40 µM GPN at 400 sec. The bar graph shows the quantification of the Δ[Ca^2+^] from three biological replicates. Data are presented as mean ± SD. p value: unpaired two tailed t test performed in all comparisons.

Most importantly, MgATP-driven Ca^2+^ uptake by permeabilized cells measured using Mag-Fluo-4, showed no TgSERCA activity after 48 h of culture with ATc (**Fig. 4C**). This experiment validated the Mag-Fluo-4 method for assessing SERCA activity. At 24 hours post-culture with ATc, some residual SERCA activity was still detected (**Fig. 4C**).

Interestingly, Ca^2+^ entry measured in Fura-2 loaded *i!ιTgSERCA* parasites (±ATc) was not affected by the downregulation of TgSERCA (**Fig. 4D**). This finding argues against the presence of an ER-based mechanism that regulates Ca^2+^ entry. Moreover, the cytosolic resting Ca^2+^ concentration remained unchanged in the *i!ιTgSERCA* (+ATc) mutant (**Fig. 4** **A,B, D-H**) highlighting a critical role of the plasma membrane Ca²⁺ pump in maintaining cytosolic Ca²⁺ homeostasis.

The response to Zaprinast was diminished but was still present (**Fig. 4E**) indicating that Zaprinast induces Ca^2+^ release from the ER and from an additional compartment. When Zaprinast was added after Ca²⁺ replenishment, the response remained reduced in the mutant pre-incubated with ATc (**Fig. 4F**). We next tested GPN, which primarily targets acidic stores, and observed a decreased response (**Fig. 4G**). Adding, GPN after replenishing the cells with Ca^2+^ resulted in an increased response as we showed in **Fig. 1**, but this response was also reduced when the mutant was grown with ATc (**Fig. 4H**).

Given that the Ca²⁺ phenotypes were assessed after 24 hours of ATc treatment, when approximately 50% of TgSERCA activity remains, the response to Zaprinast may still reflect ER involvement, and may not provide definitive evidence for the contribution of an additional Ca²⁺ pool. To further investigate this, we conducted an experiment in which TG was added prior to GPN and Zaprinast. In this setting, GPN significantly reduced the Zaprinast-induced response (**Fig. S2B**). This result suggests that Zaprinast also targets a non-ER Ca²⁺ store, and that this store is likely the same one affected by GPN.

These results support a functional link between the stores targeted by GPN and the ER. Given that SERCA downregulation impaired ER Ca²⁺ storage without affecting cytosolic Ca²⁺ uptake and cytosolic Ca^2+^ levels, the diminished response to GPN suggests that Ca²⁺ released or leaked from the ER is important for refilling the store targeted by GPN.

### The mitochondrion takes up Ca^2+^ from the ER and from acidic stores

In mammalian cells, the high concentration of Ca^2+^ in the ER is important for mitochondrial ATP production (Wenzel *et al*., 2022). This is because of the close proximity between the ER and mitochondria which allows for the directional flow of Ca^2+^ from the ER to the mitochondria (Gincel *et al*., 2001; Rapizzi *et al*., 2002). With the aim of verifying if the *T. gondii* mitochondria can take up calcium, we introduced a genetic Ca^2+^ indicator in the mitochondrion of *T. gondii* tachyzoites by attaching the *GCaMP6f* gene (Chen *et al*., 2013) to the mitochondrial targeting signal of the *T. gondii* superoxide dismutase 2 (SOD2) gene (Pino *et al*., 2007) and isolated stable transgenic clones (RH*-SOD2-GCaMP6f*) (Vella *et al*., 2020). Fluorescence microscopy of live cells confirmed GCaMP6f localization to the mitochondria (**Fig. 5A**). Direct Ca^2+^ uptake was observed in digitonin permeabilized parasites incubated in the presence of increasing concentrations of Ca^2+^ (**Fig. 5B-C**). Although a measurable increase in mitochondrial fluorescence was observed, it required high Ca²⁺ concentrations, indicating that the *T. gondii* mitochondrion can take up Ca²⁺ but do so with very low affinity. These Ca²⁺ levels were significantly higher than the typical cytosolic Ca²⁺ concentrations found in healthy cells (**Fig. 5B-C**).

**Figure 5:**
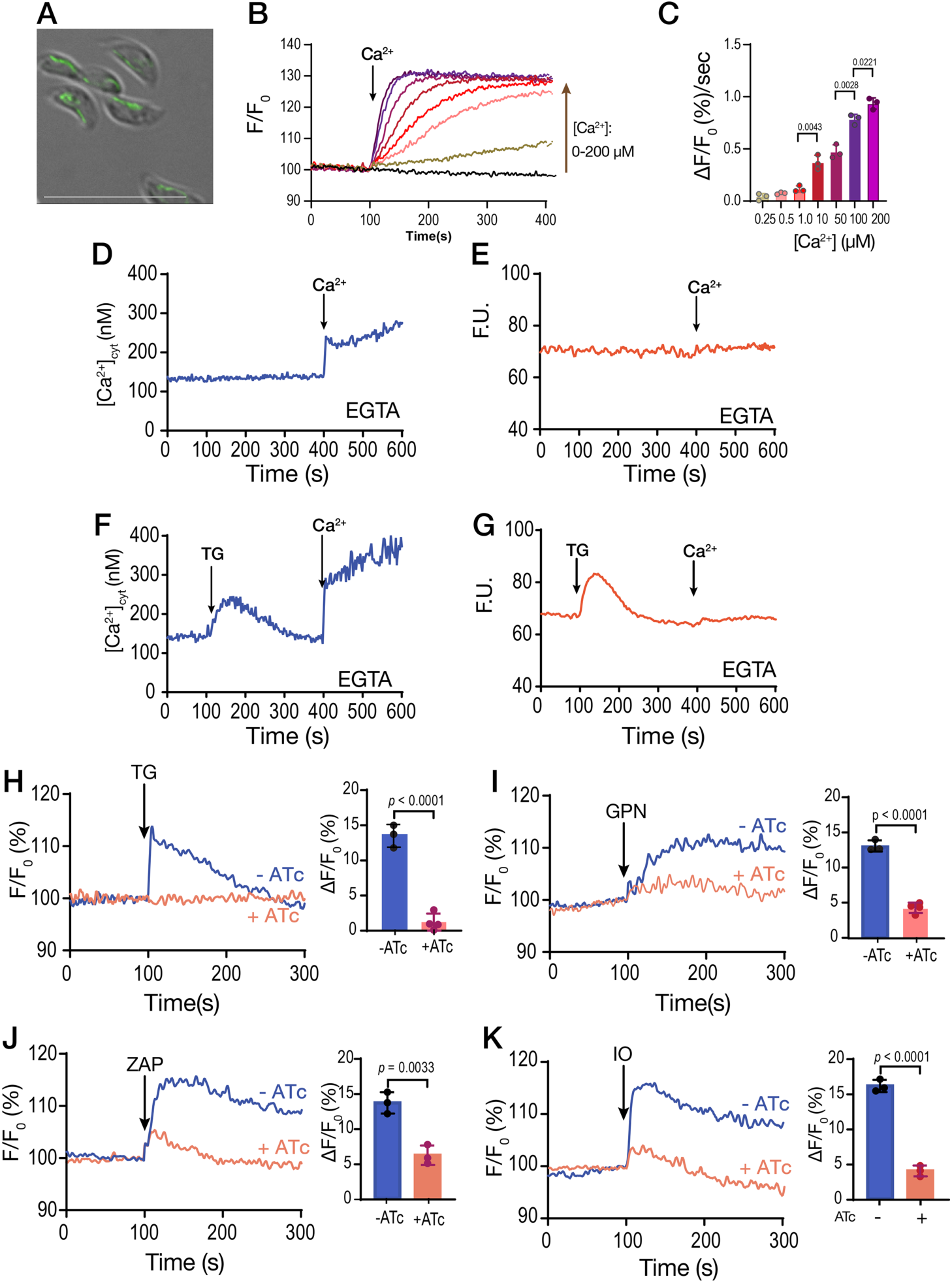
**Mitochondrial Calcium uptake**. **A**, Fluorescence image of *T. gondii* tachyzoites of the RH strain expressing *SOD2-GCaMP6f* (pDT7S4H3-SOD2-GCaMP6f). The generation of this cell line is described in the methods section. **B**, Ca²⁺ uptake in digitonin-permeabilized *T. gondii* tachyzoites expressing SOD2-GCaMP6f. Parasites (5 × 10⁷) were permeabilized as described in the Methods section and suspended in buffer containing 100 µM EGTA. Ca²⁺ was added at 100 s to reach final free concentrations of 0.25, 0.5, 1, 10, 50, 100, and 200 µM, calculated using Maxchelator. **C**, ΔF was measured as the change in fluorescence between the baseline and the maximum value obtained 20 s after Ca²⁺ addition. Data represent the average of three independent biological experiments. **D**, Fura-2-loaded *T. gondii* tachyzoites expressing SOD2-GCaMP6f in suspension. The experimental setup was identical to that described in Fig. 1A-B. CaCl₂ (1.8 mM) was added at 400 s, and fluorescence measurements were performed under Fura-2 conditions. **E**, GCaMP6f fluorescence measurements of intact *T. gondii* tachyzoites expressing SOD2-GCaMP6f targeted to the mitochondrion. Fluorescence was recorded using optimized settings for GCaMP6 detection. **F**, 1 µM TG was added at 100 sec followed by 1.8 mM CaCl2 at 400 sec. Fura-2 loaded parasites and Fura-2 conditions were used. **G**, Same additions as in F but measuring fluorescence of GCaMP6f. **H**, Response to 1 µM TG of *iΔTgSERCA*-SOD2-GCaMP6f parasites (transfected with the pCTH3-SOD2-GCaMP6f plasmid), grown with (pink trace) or without (blue trace) ATc. Fluorescence measurements were performed under the same conditions as in panel G using intact parasites. The bar graph shows ΔF values from three independent biological replicates. **I**, same as H but using 40 μM GPN. **J**, Same as H but using 100 μM Zaprinast. **K**, Same as H but using 1 μM Ionomycin. Data are presented as mean ± SD from 3 independent biological experiments. *p value*: unpaired two tailed t test performed in all comparisons.

We hypothesized that the *T. gondii* mitochondrion may take up Ca²⁺ through close membrane contacts with the ER, where localized Ca²⁺ concentrations in microdomains could be significantly higher than in the cytosol, a mechanism previously described in mammalian cells (Rizzuto *et al*., 1998). We next loaded the RH*-SOD2-GCaMP6f* mutant with Fura-2 to simultaneously monitor cytosolic and mitochondrial Ca²⁺ in intact parasites. Upon addition of extracellular Ca²⁺, an increase in cytosolic Ca²⁺ was observed, however mitochondrial GCaMP6f fluorescence remained unchanged (**Fig. 5D-E**), suggesting that mitochondria are unable to take up Ca²⁺ at the cytosolic concentrations reached under these conditions. This also validates the proper localization of the indicator, confirming its absence from the cytosol. Addition of TG followed by extracellular Ca²⁺ resulted in a cytosolic Ca²⁺ increase, readily detected in Fura-2-loaded parasites. However, and most importantly, only TG triggered a measurable increase in the mitochondrial GCaMP6f signal, whereas a rise in cytosolic Ca²⁺ induced by extracellular Ca²⁺ addition alone did not (**Fig. 5G**). Our interpretation is that TG-induced ER Ca²⁺ leakage led to local accumulation of Ca²⁺ at the cytosolic face of the ER membrane, creating microdomains of high Ca²⁺ concentration sufficient to trigger mitochondrial uptake. Addition of Ca²⁺ after TG resulted in a greater increase in cytosolic Ca²⁺ (**Fig. 5F**) compared to TG alone. However, even under these conditions, no corresponding increase in mitochondrial GCaMP6f fluorescence was observed. This further confirms that mitochondria are unable to take up cytosolic Ca²⁺ at these low concentrations.

We next introduced the same *SOD2*-GCaMP6f chimeric gene into the *iΔTgSERCA* mutant background and isolated a clonal line (*iΔTgSERCA-SOD2-GCaMP6f*) (**Fig. S5A-B**). Fluorescence imaging of live cells confirmed proper localization of the indicator (**Fig. S5A**), and fluorescence measurements of intact cells corroborated that adding extracellular Ca²⁺ did not increase GCaMP6f fluorescence, whereas addition of TG to the suspension resulted in a fluorescence increase (**Fig. S5B**). We next monitored changes in GCaMP6f fluorescence in the mutant and compared results between parasites grown with and without ATc. In line with prior observations, parasites cultured without ATc showed a consistent and measurable increase in mitochondrial GCaMP6f signal upon TG treatment (**Fig. 5H**, *blue trace*). In contrast, this response was abolished in parasites cultured with ATc, consistent with reduced TgSERCA expression leading to ER Ca²⁺ depletion (**Fig. 5H**, *pink trace*).

We next tested additional stimuli and observed a clear increase in mitochondrial GCaMP6f fluorescence in the *iΔTgSERCA* mutant following the addition of GPN, Zaprinast, or IO (**Fig. 5H-J,** *blue traces*). In all cases, this fluorescence increase was significantly reduced in the *iΔTgSERCA* (+ATc) mutant (**Fig. 5H-J**, *pink traces*). These results suggest a potential direct interaction between the mitochondrion and acidic Ca²⁺ stores, such as the PLVAC and/or Golgi apparatus. The reduced Ca²⁺ content of these compartments, resulting from TgSERCA downregulation, appears to impact mitochondrial Ca²⁺ uptake.

In summary, we demonstrated that the *T. gondii* mitochondrion is capable of Ca²⁺ uptake via transfer from the ER, a process that becomes apparent upon inhibition of ER Ca²⁺ uptake with TG. This suggests that the high Ca²⁺ concentrations required for mitochondrial uptake are achieved only at membrane contact sites between the ER and mitochondria. In the *iΔTgSERCA* (+ATc) mutant, impaired ER Ca²⁺ storage due to TgSERCA downregulation compromises mitochondrial Ca²⁺ uptake. Additionally, our data suggest that the *T. gondii* mitochondrion may also take up Ca²⁺ from acidic stores, such as the PLVAC or Golgi, which appear to rely indirectly on ER Ca²⁺ refilling. When TgSERCA is downregulated, depletion of ER Ca²⁺ likely compromises the Ca²⁺ content of these acidic compartments, and impairing mitochondrial Ca²⁺ uptake from these stores.

### Proximity between the ER, Mitochondrion and Acidic Compartment

We next investigated if it was possible to see proximity between the ER with other organelles by IFA and/or electron microscopy (EM). We performed IFAs with ER and mitochondria markers and ER and PLVAC markers (**Fig. 6**). In intracellular parasites, the mitochondrion was observed to surround the ER, forming multiple potential sites of interaction (**Fig. 6A** and *supplemental video 1*). As described, the mitochondrion of intracellular *T. gondii* tachyzoites surrounds the periphery of the cell in a lasso-shape morphology (Ovciarikova *et al*., 2017). In contrast, in extracellular parasites, the mitochondrion displays a marked morphological change, adopting either a sperm-like or collapsed conformation (Ovciarikova *et al*., 2017) (**Fig. 6B** and *supplemental video 2*). Our hypothesis is that this retraction of the mitochondrion allows the ER membranes to expand in extracellular parasites and extend toward the apical domain of the parasite where Ca^2+^ is needed for secretion of micronemes and conoid extrusion.

**Figure 6:**
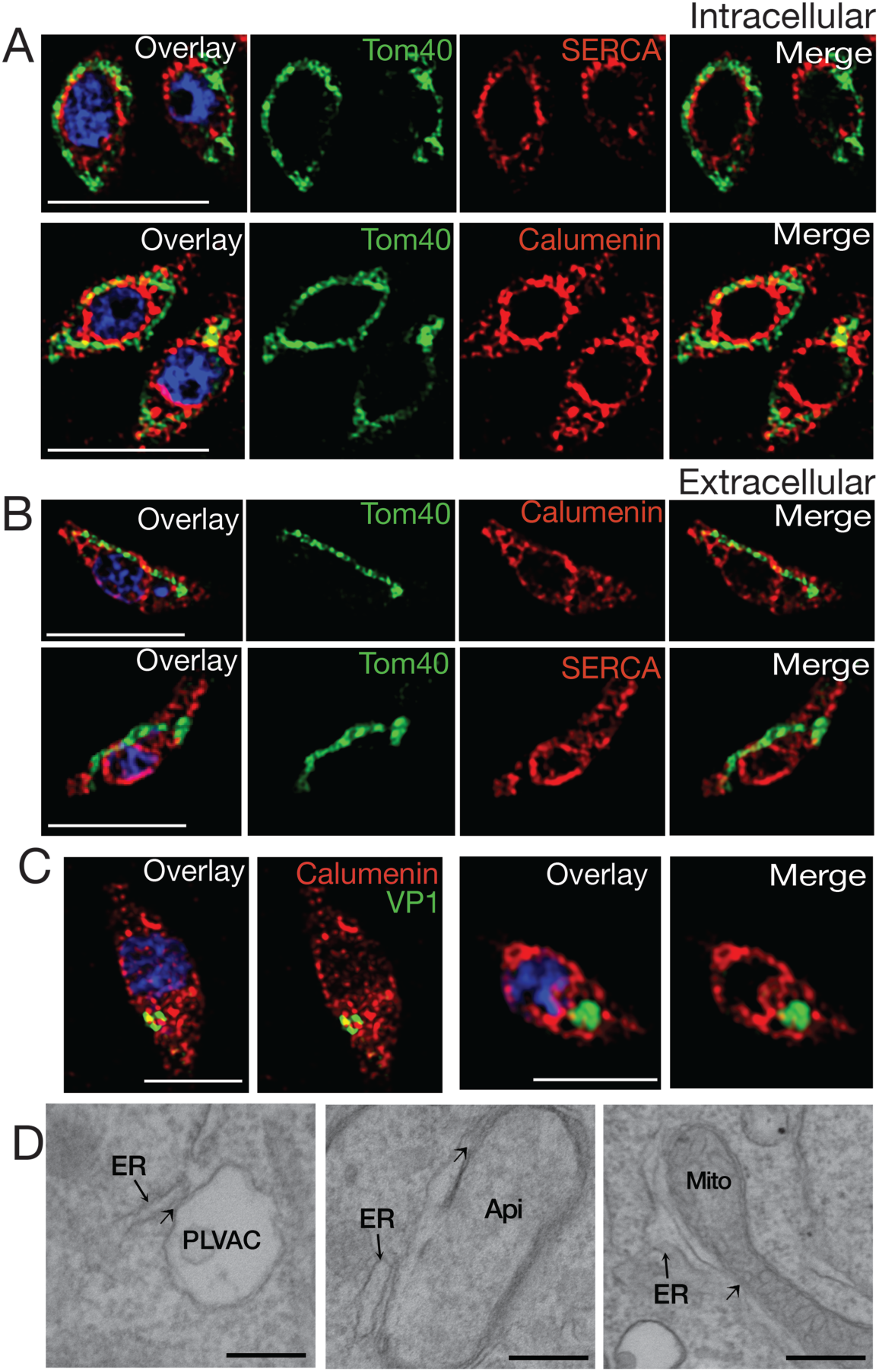
ER membrane contacts with the mitochondrion and the plant like vacuolar compartment (PLVAC). **A**, Super-resolution IFAs of intracellular parasites with the mitochondrion labeled with the αTom40 (green, 1:20,000) antibody and the ER labeled with the αTgcalumenin antibody (an ER calcium binding protein) (red, 1:1000) or the TgSERCA (red 1:1000). **B**, IFAs of extracellular tachyzoites with the same antibodies used for part A. The mitochondrion and ER membranes interact in several regions. **C**, The PLVAC was labeled with the αTgVP1 antibody (green, 1:200) or the αTgCPL antibody (green, 1:500). The ER was labeled with the αTgcalumenin antibody (red). The points of contact between the ER and the PLVAC are yellow. Scale bars in A-C is 5 µm. **D**, Transmission Electron Microscopy imaging of the contact sites formed between ER and PLVAC, ER and Apicoplast, ER and mitochondria. Size bars are 100 nm.

The PLVAC also formed points of contact with the ER (**Fig. 6C** and *supplemental video 3*). Multiple points of contact were also observed by EM between the ER and the PLVAC, the ER and the apicoplast and the ER and the mitochondrion (**Fig. 6D**).

Interestingly these contacts were still present in the *i!ιTgSERCA* (+ATc) mutant (**Fig. S6, A-D**), as most likely TgSERCA would not be directly involved in the establishment of contacts. We quantified the length of the limiting membrane of the organelle in contact with ER membranes at a distance of less than 30 nM and found that after knockdown of TgSERCA, the contact was not altered. However, Ca^2+^ transported from the ER into the mitochondrion after TG treatment was significantly decreased which means that this phenotype is due to reduced ER Ca^2+^ and not to lack of contacts (**Fig. S6E**). A similar result was seen when measuring contacts between the ER and the apicoplast (**Fig. S6F**) and between the ER and the PLVAC (**Fig. S6G**).

In summary, this data supports the presence of points of contact between the ER and other organelles like the mitochondrion, the PLVAC and the apicoplast. These contacts likely facilitate the transfer of Ca²⁺ from the ER, the organelle with the highest Ca²⁺ content, to other compartments.

## DISCUSSION

In this work, we demonstrated that the endoplasmic reticulum (ER) of *T. gondii* has a remarkable capacity to sequester Ca²⁺ entering the cytosol from the extracellular milieu, achieving this with only a minimal rise in cytosolic Ca²⁺ levels. This is largely due to the activity of a highly efficient SERCA Ca^2+-^ATPase (TgSERCA), which has a high affinity for Ca^2+^. The activity of TgSERCA, most likely together with the plasma membrane Ca²⁺ pump (Luo et al., 2001; Luo et al., 2005), limits large increases in cytosolic Ca²⁺ (Hortua Triana et al., 2024).

We provide evidence that the ER not only sequesters extracellular Ca²⁺ through TgSERCA activity but also shares this pool with other organelles, including mitochondria and acidic stores. This capacity stems from the unique ability of the ER to capture a sizable fraction of extracellular Ca²⁺ entering the tachyzoite cytosol. Such inter-organelle transfer allows localized Ca²⁺ release without globally elevating cytosolic levels, thereby preventing unintended signaling events. Our data support a model in which loss of SERCA activity reduces ER Ca²⁺ as well as Ca²⁺ content in other organelles. Under physiological conditions, ER Ca²⁺ is regularly mobilized for signaling and homeostasis, helping to maintain Ca²⁺ balance across cellular compartments (see our hypothetical model in Fig. 7).

**Figure 7:**
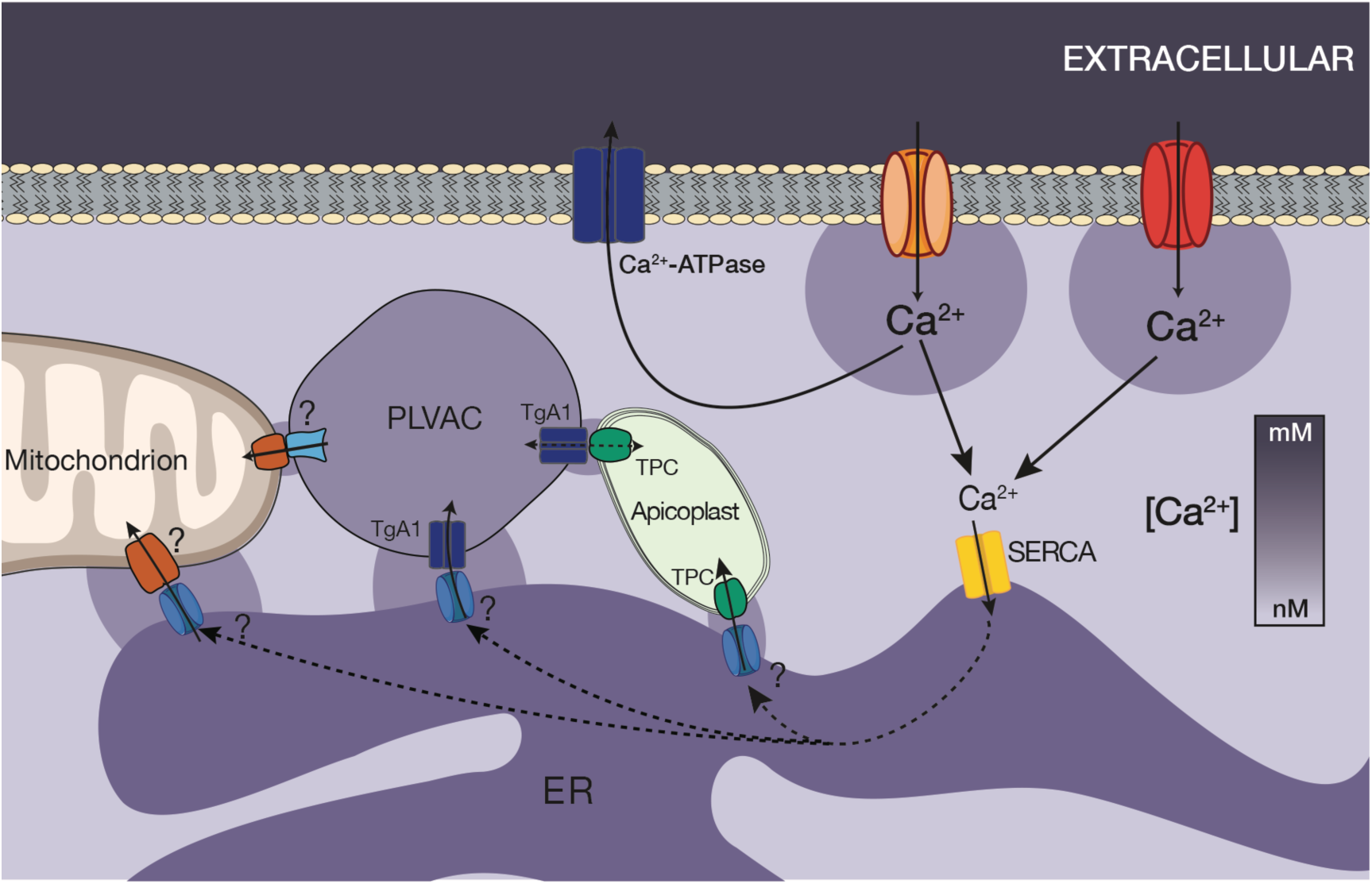
Hypothetical model showing calcium entry through two different types of Ca^2+^ channels, uptake by TgSERCA into the ER and distribution to the other organelles via transfer from the ER to the mitochondria, PLVAC and apicoplast. The mitochondrion is shown in close contact to the ER which constitutively leaks Ca^2+^ into the cytosol. Ca^2+^ could leak from the ER through the TgTRPPL-2 channel previously described (Marquez- Nogueras et al., 2021). The mitochondria take up Ca^2+^ from the ER through an unknown mechanism. VDAC could be involved in the transfer through the outer mitochondrial membrane (Mallo et al., 2021). The PLVAC interacts with the ER and may also interact with the mitochondrion and the apicoplast. TgA1, a calcium ATPase previously characterized may be the pump involved in Ca^2+^ uptake (Luo et al., 2001; Luo et al., 2005). The mechanism of release is unknown. The Two Pore Channel (TgTPC) was shown to be involved in the formation of contacts between the ER and the apicoplast (Li et al., 2021) and could form part of the mechanism of uptake. Question marks point to molecules or mechanism partially or not yet identified.

SERCA Ca²⁺-ATPases are P-type pumps located in the ER and secretory pathway membranes (Kuhlbrandt, 2004). Mammals express three isoforms (SERCA1–3), with SERCA2b serving as the housekeeping form (Wuytack *et al*., 2002). SERCA pumps translocate two Ca²⁺ ions into the ER lumen per ATP hydrolyzed, lowering cytosolic Ca²⁺ to resting levels (<100 nM) and replenishing ER stores (∼500 µM). This stored Ca²⁺ supports signaling and the activity of luminal enzymes critical for cell growth, proliferation, and differentiation (Wuytack *et al*., 2002).

*T. gondii* appears to express a single SERCA protein (TgSERCA) (Nagamune and Sibley, 2006), likely serving a housekeeping role. The severe defects observed in the *iΔTgSERCA* (+ATc) mutant like impaired replication and disruption of the lytic cycle, highlight its essential function. TgSERCA activity was dependent on MgATP and exhibited high Ca²⁺ affinity, as evidenced by Mag-Fluo-4-based assays detecting uptake at free Ca²⁺ levels as low as 55 nM. This suggests that TgSERCA functions effectively at physiological cytosolic Ca²⁺ concentrations (60–100 nM), ensuring ER loading even under resting conditions.

*In situ* characterization of organellar Ca²⁺ uptake has been feasible in trypanosomes (Docampo and Vercesi, 1989; Vercesi *et al*., 1990) but remains challenging in *T. gondii*. The Mag-Fluo-4 and the mitochondrial GCaMP6f protocols enable reliable measurement of ER and mitochondrial Ca²⁺ uptake, respectively. For ER Ca^2+^ uptake MgATP was essential for the activity of TgSERCA, as other forms of ATP were ineffective. This protocol has been extensively used in mammalian cells, DT40 cells and other cells for measuring intraluminal calcium, activity of SERCA and response to IP3 (Laude *et al*., 2005; Rossi *et al*., 2009; Valverde *et al*., 2010; Sampieri *et al*., 2018; Rossi and Taylor, 2020). We previously successfully employed it for the characterization of the *Trypanosoma brucei* IP3R (Huang *et al*., 2013) and the assessment of SERCA activity in *T. gondii* mutants (Li *et al*., 2021). In this work, we used it to assess TgSERCA activity under defined Ca²⁺ and MgATP conditions.

Using the Mag-Fluo-4 protocol, we observed that cyclopiazonic acid (CPA), a reversible SERCA inhibitor (Inesi and Sagara, 1994), induced a Ca^2+^ leak rate comparable to the one after adding TG. This indicates that the leak rate is mainly determined by intrinsic leak mechanisms rather than the type of SERCA inhibition. However, in intact Fura-2-loaded parasites, CPA induced a smaller cytosolic Ca²⁺ increase than TG. This likely reflects CPA’s reversible and potentially incomplete inhibition of SERCA under cellular conditions, as was also observed in *Plasmodium falciparum* (Borges-Pereira *et al*., 2020).

We observed that the response to acidic calcium triggers like nigericin or GPN were greatly enhanced when added after TG in Fura-2-loaded tachyzoites, likely due to ER Ca²⁺ leak and subsequent transfer to other compartments. Additionally, downregulation of *TgSERCA* expression resulted in reduced responses to these acidic store triggers, supporting the notion that the ER contributes to the filling of these organelles. It is important to note that our analyses of Ca²⁺ storage was done in parasites that retained partial TgSERCA activity, as it is not possible to isolate cells entirely lacking TgSERCA expression. Under these conditions, Zaprinast still induced a reduced Ca²⁺ mobilization response. This residual response may be due to remaining calcium in the ER or may suggest that Zaprinast targets multiple calcium stores. We recently identified the Golgi apparatus as a calcium store in *T. gondii* (Calixto *et al*., 2025) and demonstrated that treatment with GPN in Fura-2-loaded tachyzoites diminished the Zaprinast-induced calcium response, suggesting that Zaprinast and GPN may act on overlapping stores. In the present study, we demonstrated that sequential treatment with TG followed by GPN almost completely abolished the Zaprinast response, further supporting this idea. Although GPN is primarily known to act on acidic organelles, it has also been proposed to affect the ER (Atakpa *et al*., 2019) however, we have no evidence that GPN mobilizes calcium from the ER in *T. gondii*. We propose that GPN primarily targets the PLVAC, but further investigation is required to fully characterize its mechanism of action.

It was interesting that Ca²⁺ entry remained unchanged in the *i!ιTgSERCA* (+ATc) mutant, suggesting that intracellular stores may not be directly involved in the regulation of Ca²⁺ entry (Pace *et al*., 2014). Moreover, conventional genomic analysis has not identified homologs of the canonical Store-Operated Calcium Entry (SOCE) components, STIM and Orai (Collins and Meyer, 2011), raising the possibility that these proteins are either absent or highly divergent in sequence and lack conserved regulatory domains. If communication between intracellular stores and the plasma membrane exists in *T. gondii*, the underlying mechanism remains unclear.

Calcium transport into the *T. gondii* mitochondrion had not been previously demonstrated and our findings provide the first experimental evidence for this process, though the molecular mechanism remains unclear. We found that normal cytosolic Ca²⁺ fluctuations were insufficient to drive mitochondrial uptake, consistent with the low Ca²⁺ affinity of the mitochondrion. Uptake occurred only after SERCA inhibition, which caused local Ca²⁺ accumulation at the cytosolic side of the ER membrane, enabling transfer to the mitochondrion, likely via membrane contact sites (MCSs), since direct uptake from the cytosol would be inefficient at low Ca²⁺ concentrations. MCSs were defined as stable, tightly apposed, but non-fusogenic regions of close proximity between subcellular organelles, and play a key role in inter-organelle communication (Phillips and Voeltz, 2016). In mammalian cells, the ER forms an extensive network of MCSs with the PM, mitochondria and endocytic vesicles for the exchange of Ca^2+^ (Burgoyne *et al*., 2015).

In *T. gondii* the characterization of MCSs is only in its beginnings (Huet and Moreno, 2023) with only a few evidences for their presence (Tomova *et al*., 2009; Mallo *et al*., 2021; Ovciarikova *et al*., 2022; Ovciarikova *et al*., 2024) and function (Li *et al*., 2021; Oliveira Souza *et al*., 2022). Imaging of intracellular *T. gondii* showed that its mitochondrion surrounds the periphery of the cell in a lasso-shape conformation. On the other hand, in extracellular parasites the mitochondrion changes its morphology and adopts a sperm-like or collapsed conformation (Ovciarikova *et al*., 2017). Our IFA analysis with ER and mitochondrial markers revealed that the lasso-shaped mitochondrion surrounds the ER with plenty of opportunities for contact between both organelles. Interestingly, the mitochondrion also appeared capable of importing Ca²⁺ from acidic stores such as the PLVAC, as GPN treatment stimulated mitochondrial Ca²⁺ uptake. This response was reduced in the *iΔTgSERCA* (+ATc) mutant, indicating that TgSERCA activity contributes to this process. Moreover, ER Ca²⁺ depletion impaired transfer from acidic stores to the mitochondrion, suggesting functional interdependence among Ca²⁺ stores and a central role for the ER in coordinating intracellular Ca²⁺ dynamics.

In mammalian cells Ca^2+^ ion is transferred from the ER to the mitochondrion through the outer membrane voltage-dependent anion channel1 (VDAC1) (Gincel *et al*., 2001; Rapizzi *et al*., 2002) and the inner membrane calcium uniporter (MCU1) (De Stefani *et al*., 2011). A VDAC homologue is present in *T. gondii*, which was shown to be essential for growth and for mitochondrial and ER morphology (Mallo *et al*., 2021). However, molecular evidence for the presence of a Ca^2+^ uniporter in the inner mitochondrial membrane, driven by the electrochemical gradient generated by the electron transport chain, remains to be demonstrated.

Cellular responses triggered by Ca²⁺ signals are shaped by the location, duration, and amplitude of the signals. Movement of Ca^2+^ in the cytosol of cells is severely limited due to the presence of high affinity Ca^2+^ buffers. In mammalian cells it was shown that Ca^2+^ tunnels through the ER as it moves faster because the Ca^2+^ binding capacity of the ER is almost 100 lower than the binding capacity of the cytosol (Mogami *et al*., 1997). The ER Ca^2+^ transport through its lumen was shown to provide a mechanism of delivering Ca^2+^ to targeted sites without activating inappropriate processes in the cell cytosol (Petersen *et al*., 2017). In *T. gondii*, the relative Ca²⁺-binding capacity of the cytosol compared to the ER remains poorly understood, as the localization of many predicted Ca²⁺-binding proteins has not been fully determined. Several calmodulin-like proteins, for example, are localized to the conoid (Long *et al*., 2017).

The mechanisms of Ca²⁺ entry at the plasma membrane, release from the ER, and uptake by the mitochondria or acidic stores remain incompletely characterized (Hortua Triana *et al*., 2018; Pace *et al*., 2020) (Garcia *et al*., 2017). Consequently, the molecular elements required for classical Ca²⁺ tunneling have not been identified in *T. gondii*. Nevertheless, our results demonstrate that Ca²⁺ can be transferred from the ER to other organelles. This is supported by the increased mitochondrial and acidic calcium pools observed following pharmacological ER depletion, both in the presence and absence of extracellular calcium. Importantly, chronic ER calcium depletion, such as in the *iΔTgSERCA* mutant cultured with ATc, leads to the depletion of all intracellular Ca²⁺ stores. Additionally, we directly demonstrated mitochondrial Ca²⁺ uptake when Ca²⁺ accumulated on the cytosolic side of the ER membrane following SERCA inhibition. The specific roles of Ca²⁺ in the mitochondrion and acidic compartments remain unclear. In mitochondria, Ca²⁺ may support ATP production, although this has yet to be confirmed. Both organelles may also act as auxiliary Ca²⁺ reservoirs during ER Ca²⁺ overload.

In *T. gondii* cytoplasmic Ca^2+^ increases, due to efflux from the ER or entry through the PM, have been reported to initiate key parasite processes such as microneme secretion (Carruthers and Sibley, 1999; Nagamune *et al*., 2007b), conoid extrusion (Del Carmen *et al*., 2009) (Pace *et al*., 2014), invasion (Vieira and Moreno, 2000; Lovett and Sibley, 2003) and egress (Arrizabalaga *et al*., 2004; Borges-Pereira *et al*., 2015). These responses require precise spatiotemporal regulation of Ca²⁺ at specific cellular sites, suggesting the presence of mechanisms that direct Ca²⁺ to discrete locations. We propose that the ER plays a central role in this regulation by acting as a hub that distributes Ca²⁺ to defined sites at defined times to initiate parasite functions. The severe invasion, replication, and egress defects observed in the *iΔTgSERCA* (+ATc) mutant support this hypothesis. Egress was one of the first steps of the *T. gondii* lytic cycle that was shown to be triggered by exposure of intracellular parasites to ionophores (Endo *et al*., 1982). Most recent work using GECIs demonstrated the rise in cytosolic calcium preceding egress (Borges-Pereira *et al*., 2015; Stewart *et al*., 2017). In the present study, we demonstrate that egress is defective and unresponsive to ionophores in parasites lacking sufficient Ca^2+^ in their intracellular stores. This underscores the critical role of TgSERCA in maintaining Ca^2+^ stores filled. Interestingly, this defect could be rescued by exposing the parasites to extracellular calcium through host cell permeabilization, which not only restored egress but also was faster.

In conclusion, this study demonstrates that the ER of *T. gondii* can replenish itself with Ca^2+^ and acts as a source of Ca^2+^ for cytosolic signaling, as well as for loading acidic stores and the mitochondrion. *T. gondii* is a protozoan parasite that causes disease by reiterating a lytic cycle that is driven by Ca^2+^ signaling. Our findings enhance understanding of how extracellular and intracellular Ca^2+^ stores coordinate to sustain the pathologic features of *T. gondii*. Future studies will focus on defining the roles of Ca^2+^ in mitochondrial and acidic stores functions.

## Methods

### Cell Culture

*Toxoplasma gondii* tachyzoites (RH and *TatiΔku80* strain) were maintained in human telomerase reverse transcriptase immortalized foreskin fibroblasts (hTERT) (Farwell *et al*., 2000) grown in Dulbecco’s modified minimal essential media (DMEM) with 1% FBS. These cells are tested for Mycoplasma contamination regularly and are treated with mycoplasma removal agent.

### Generation of SERCA mutants

A promoter insertion plasmid was generated by cloning three PCR fragments into a modified pCR2.1-TOPO vector using Gibson Assembly Cloning Kit (NEB #E5510). One fragment corresponding to the TgSERCA flanking region (predicted promoter/5’UTR) was amplified with primers 1 and 2 (Table S1). The second fragment corresponds to DHFR+T7S4 (Sheiner *et al*., 2011) and was amplified with primers 3 and 4. Another fragment corresponds to the 5’ TgSERCA coding sequence beginning with start codon and was amplified with primers 5 and 6. The vector pCR2.1-TOPO was used, which had only one EcoRI site and it was cut with the enzyme NotI to use as a vector backbone. The promoter insertion plasmid was transfected into the *TatiΔku80* cells and selected with 1 μM pyrimethamine using a “ultra aggressive” screening method. Briefly, 200 μl of the suspension of transfected parasites was added to 10 ml of media and then one to three drops (∼65 μl per drop) were inoculated into three 24-well plates containing media. The clonal lines created after selection and subcloning were termed *iΔTgSERCA*.

For *in situ* tagging, an approximately 2 kb fragment was amplified from the genomic locus (3’ region) of the TgSERCA gene using primers 7 and 8. The fragment was cloned into the pLic-3HA- CAT plasmid (Huynh and Carruthers, 2009) and the construct was linearized with the enzyme NheI for transfection into the *iΔTgSERCA* mutant. Clonal cell lines were generated after selection with chloramphenicol and subcloning and termed *iΔTgSERCA-3HA*.

### Expression and purification of TgSERCA recombinant protein

The phosphorylation (P) and nucleotide binding(N) domains of *TgSERCA* (TGGT1_230420) (nucleotides 1123 to 2415, amino acid residues 375 to 805) were cloned into XmaI and HindIII sites of pQE-80L with primers 13 and 14 to create recombinant protein with a N-terminal 6xHis tag. The resulting plasmid was transformed into *Escherichia coli* BL21-CodonPlus competent cells and expression was induced by addition of 0.4 mM isopropyl Δ-D-1-thiogalactopyranoside (IPTG) for 4 hours at 37°C. Cells were pelleted and resuspended in equilibration/binding buffer (50 mM Na3PO4, 300 mM NaCl, 10 mM Imidazole, 8 M Urea, and protease inhibitor cocktail P-8849). The cells were then sonicated for 80 seconds and centrifuged at 12,000 rpm for 20 min at 4°C. The supernatant was filtered through a 0.45 μm membrane and the protein was purified using HisPur Ni-NTA Chromatography Cartridge (Thermo Scientific) following instructions from the manufacturer. Proteins that were unbound were washed with 12 ml of wash buffer (50 mM Na3PO4, 300 mM NaCl, 40 mM imidizole, and 8 M urea), and the recombinant protein was eluted with 5 ml elution buffer (50 mM Na3PO4, 300 mM NaCl, 250 mM imidazole, and 8 M urea). Eluted protein fractions were concentrated and desalted with Amicon Ultra-0.5 mL centrifugal filter (Millipore Sigma).

### Anti-TgSERCA antibody generation in Guinea pigs

Two Guinea pigs were each immunized with 0.2 mg of purified recombinant protein mixed with equal volume of Freund’s Complete Adjuvant (Sigma F5581), followed by two boosts of 0.1 mg antigen mixed with equal volume of Freund’s Incomplete Adjuvant (Sigma F5506) for Guinea pig 1 and three boosts for Guinea pig 2. The resulting antibodies were tested at 1:1,000 in western blot against RH lysates and were developed with Alexa Fluor 488 goat anti-Guinea pig (1:1,000). The antibodies were compared with Dr. Sibley’s mouse anti-SERCA antibody (Nagamune *et al*., 2007a) to confirm size and purity. The anti-TgSERCA antibodies were then affinity purified. Guinea pigs were handled according to our approved institutional animal care and use committee (IACUC) protocols (A2021 03-005-A5) of the University of Georgia.

### Cytosolic calcium measurements with Fura-2

*T. gondii* tachyzoites were loaded with Fura-2 AM as previously described (Vella *et al*., 2020; Stasic *et al*., 2021). Freshly released tachyzoites were washed twice with buffer A plus glucose (BAG; 116 mM NaCl, 5.4 mM KCl, 0.8 mM MgSO_4_, 50 mM HEPES, pH 7.3, and 5.5 mM glucose), by centrifugation (706 x *g* for 10 min) and re-suspended to a final density of 1 x l0^9^ parasites/ml in loading buffer (BAG plus 1.5% sucrose, and 5 μM Fura-2-AM). The suspension was incubated for 26 min at 26 °C with mild agitation. Subsequently, the parasites were washed twice (2,000 x *g* for 2 min) with BAG to remove extracellular dye, re-suspended to a final density of 1x10^9^ parasites per ml in BAG and kept on ice. This loading protocol is specifically designed to minimize Fura-2 compartmentalization, which is typically indicated by elevated resting Ca²⁺ concentrations. All experiments are conducted within a time frame during which resting Ca²⁺ levels remain stable, typically below or at 100 nM. For fluorescence measurements, 2 x 10^7^ parasites/mL were placed in a cuvette with 2.5 mL of Ringer’s buffer without calcium (155 mM NaCl, 3 mM KCl, 1 mM MgCl2, 3 mM NaH2PO4, and 10 mM Hepes, and 10 mM glucose). Fluorescence measurements were done in a Hitachi F-7000 or F-4500 fluorescence spectrophotometer using the Fura-2 conditions for excitation (340 and 380 nm) and emission (510 nm). The Fura-2 fluorescence response to Ca^2+^ was calibrated from the ratio of 340/380 nm fluorescence values after subtraction of the background fluorescence of the cells at 340 and 380 nm as previously described (Grynkiewicz *et al*., 1985). The Ca^2+^ release rate was defined as the change in Ca^2+^ concentration during the initial 20 s after reagent addition. Delta F was calculated by the difference between the higher calcium peak and basal Ca^2+^, and recovery was defined as the change of Ca^2+^ concentration after the calcium peak was reached for the subsequent indicated times.

### Endoplasmic reticulum Ca^2+^ measurements in permeabilized T. gondii tachyzoites

Tachyzoites freshly egressed and washed as described above were resuspend to a final density of 1x10^9^ cells/ml in HBS buffer (135 mM NaCl, 5.9 mM KCl, 1.2 mM MgCl2, 11.6 mM HEPES pH 7.3, 1.5 mM CaCl2, 11.5 mM glucose) containing 1 mg/ml BSA, 0.2 mg/ml of pluronic F127 and 20 μM Mag-Fluo4-AM. The suspension was incubated at RT with mild shaking for 1 h, in the dark. Subsequently, parasites were washed 2 times at 5,000 rpm for 2 min to remove extracellular dye. The pellet was resuspended in 1.8 ml of CLM buffer (20 mM NaCl, 140 mM KCl, 20 mM PIPES, pH 7.0) containing 1 mM EGTA at 1x10^9^ cells/ml. Cells were permeabilized with 44.4 μg/ml digitonin for 6 min. Cells were washed 2 times with CLM containing 1 mM EGTA at 5,000 rpm for 2 min to remove digitonin and resuspended to a final density of 1x10^9^ tachyzoites/ml and kept in ice. For each test, 50 μl (5x10^7^) of parasite suspension was added to 1.95 ml of CLM containing 1 mM EGTA and 0.375 mM CaCl2 which results in 220 nM free Ca^2+^ as calculated with MaxChelator. Fluorescence was measured with a Hitachi F-7000 or F-4500 fluorescence spectrophotometer (Excitation at 485 nm and emission at 520 nm). Ratio (ΔF/F0/sec) was evaluated by measuring the rate of change in fluorescence over 20 s after reagent addition.

### Strain construction and maintenance

The organelle targeting of GCaMP6f was made by overlapping PCR. The N-terminal mitochondrial targeting sequence of the *T. gondii* SOD2 gene (Pino *et al*., 2007) was used to target GCaMP6f to the mitochondrion. The *GCaMP6f* gene for this construct was amplified by primers 9 and 10 (Table S1). After gel purification of the GCaMP6f and SOD2 sequences, the mitochondria targeting construct was built by overlapping PCR with the purified PCR products as template. This construct was then cloned into the Topo-blunt vector. After the sequence was verified by sequencing, the *SOD2- GCaMP6f* fragment was removed by BglII and AvrII digestion and cloned into the same restriction sites of the pDT7S4H3 (Sheiner *et al*., 2011) and pCTH3 (Vella *et al*., 2020) vectors. The pDT7S4H3-SOD2-GCaMP6f construct was introduced into RH parasites by electroporation. After selection with pyrimethamine, the parasites were sorted by FACS and then subcloned. Clones were selected based on the dynamic range of the response to ionomycin. The pCTH3-SOD2-GCaMP6f was introduced into the *i!ιTgSERCA* mutant by electroporation. After selection with chloramphenicol, the parasites were sorted by FACS and then subcloned. The expression of GCaMP6f was verified by live cell imaging, IFA, and western blots. The clone with the largest dynamic range, as evaluated with Ionomycin, was selected to further use. **Fig. S5** shows IFA and fluorescence controls for mitochondria localization and function of the mitochondrial GCaMP6f.

### GCaMP6f fluorescence measurements

Permeabilized tachyzoites: *T. gondii* tachyzoites expressing SOD2-GCaMP6f were collected and washed 2 times at 5,000 rpm for 2 min with BAG. The parasite pellet was resuspended in 1.8 ml of BAG buffer containing 0.1 mM EGTA at 1x10^9^ cells/ml. Permeabilization with 44.4 μg/ml digitonin for 6 min was done by following the fluorescence of GCaMP6f. Cells were washed 2 times with the same buffer at 5,000 rpm for 2 min to remove digitonin, resuspended to a final density of 1 x10^9^ parasites/ml in intracellular buffer (140 mM Kgluconate, 10 mM NaCl, 2.7 mM MgSO4, 200 μM EGTA, 65 μM CaCl2, 10 mM HEPES, 10 mM Tris, pH 7.3, 1 mM Glucose) and kept in ice. 50 μl (5x10^7^) of the parasite suspension was mixed with 1.95 ml intracellular buffer for measurement. Measurements were done in a Hitachi 7000 fluorescence spectrophotometer set at 485 nm excitation and 509 nm emission. The uptake rate (ΔF/F0/Sec) was evaluated by measuring the % of change in fluorescence per second during the initial 20 s after reagent addition.

For measurements with intact parasites, they were collected, washed, and resuspended in BAG at 1x10^9^ cells/ml for testing. Fifty μl (5x10^7^) of the parasite suspension was mixed with 1.95 ml BAG containing 0.1 mM EGTA for measurement. The ratio (ΔF/F0) was evaluated by measuring the maximum change in fluorescence over 20 s after reagent addition (linear rate).

#### Growth, invasion and egress assays

Red-green invasion assays were performed as originally described (Kafsack *et al*., 2004), modified (Chasen *et al*., 2017) and adapted to use td-RFP expressing parasites. The number of tachyzoites used was 2 x 10^7^, and invasion was for 5 min. Plaque assays were performed as previously described (Roos *et al*., 1994; Liu *et al*., 2014) with modifications (Liu *et al*., 2014). 125 tachyzoites were used for infection of confluent six well plates with hTERT fibroblasts followed by an incubation time of 10 days prior to fixing and staining with crystal violet.

For egress assays, the monolayers of hTERT cells grown in 35 mm Mattek dishes were infected with 50,000 tdTomato-expressing parasites for 24 or 48 h. Parasitophorous vacuoles containing 4- 8 parasites were observed by microscopy after washing twice with Ringer’s buffer without calcium. The dish was filled with 1 ml of Ringer’s buffer supplemented with either 100 μM EGTA or 1.8 mM CaCl2. Images were collected in time-lapse mode with an acquisition rate of 3 seconds for 15 minutes. We observed that most of the *i!ιTgSERCA* cells +/- ATc were still able to egress when stimulated with 1 or 0.5 µM ionomycin added 2 min after the start of the recordings with either 100 μM EGTA or 1.8 mM CaCl2. We next used lower concentrations of ionomycin (100 nM and 50 nM) in Ringer’s buffer with 1.8 mM CaCl2 or 0.01% saponin. For egress triggered by ionomycin, the percentage of vacuoles egressed after adding ionomycin during the 15 minutes of the video (2 minutes baseline + 13 minutes after adding ionomycin) was quantified (from 100 vacuoles). For egress triggered by saponin, the time to egress after adding saponin was quantified. For natural egress, the *i!ιTgSERCA* mutant expressing td-tomato RFP was used to infect confluent hTERT cell monolayers 36 hours before adding ATc and 1 μM compound 1 (pyrrole 4- [2-(4-fluorophenyl)-5-(1-methylpiperidine-4-yl)-1H-pyrrol-3-yl]pyridine) (Cpd1) (Donald *et al*., 2002) dissolved in ethanol and the culture continued for 24 hours. After treating with Cpd1 for 24 hours, cultures showed full vacuoles which differed from the vehicle treated plates (36 hours cultured plus ATc treatment for 24 hours or 48 hours without ATc), which were fully lysed. Following synchronization, the Cpd1-containing media was removed, and the vacuoles were washed twice with warm media lacking Cpd1. Fresh media without Cpd1 was added, and the plates were transferred to a prewarmed DeltaVision microscope stage set to 37 °C. After 10 min at 37°C, egress of the full vacuoles was enumerated. We counted each plate for 1 min and we counted at least 100 vacuoles per experiment. Three independent biological experiments were conducted and summarized.

For replication assays, hTERT cells were grown on 35 mm Mattek dishes. Each dish was infected with 50,000 tdTomato-expressing parasites. 24 h after the infection counting of parasites per PV was done in a fluorescence microscope. For each experiment, at least 100 PVs were counted. Results were the average of 3 independents experiments (Li *et al*., 2021).

### Microscopy and western blot analyses

Tachyzoites were grown on hTERT cells on cover slips for ∼24 hr, washed twice with BAG and fixed with 4% formaldehyde for 1 h, followed by permeabilization with 0.3% Triton X-100 for 20 min, and blocking with 3% bovine serum albumin. IFAs were performed as previously described (Miranda *et al*., 2010). Fluorescence images were collected with an Olympus IX-71 inverted fluorescence microscope with a Photometrix CoolSnapHQ CCD camera driven by DeltaVision software (Applied Precision, Seattle, WA). Super-resolution microscopy was performed using a Zeiss ELYRA S1 (SR-SIM) system mounted on a high-resolution Axio Observer Z1 inverted microscope. The setup included transmitted light (HAL), UV (HBO), and high-power solid-state laser illumination sources (405/488/561 nm), a 100× oil immersion objective, and an Andor iXon EM-CCD camera. Image acquisition and structured illumination analysis were conducted using ZEN software (Zeiss) with the SIM analysis module. Rat anti-HA antibody (Roche) was used at a 1:25 dilution, and mouse anti-HA antibody (Covance) was used at a 1:200 dilution. Affinity-purified guinea pig anti-TgSERCA antibody was used at a 1:500 dilution.

Western blot analysis was performed as previously described (Liu *et al*., 2014). Rat anti-HA antibody from Roche was used at a dilution of 1:200. Mouse anti-HA antibody from Covance was used at a dilution of 1:1,000. The guinea pig anti-TgSERCA antibody was used at a dilution of 1:2,000. Secondary goat anti-rat or mouse antibody conjugated with HRP was used at 1:5,000. Mouse anti-α-tubulin at a dilution of 1:5,000 was used for loading control.

### Transmission Electron Microscopy

For ultrastructural observations of intracellular *T. gondii* by thin-section transmission electron microscopy (EM), infected human foreskin fibroblast cells were fixed in 2.5% glutaraldehyde in 0.1 mM sodium cacodylate (EMS) and processed as described (Coppens and Joiner, 2003). Ultrathin sections of infected host cells were stained before examination with a Hitachi 7600 EM under 80 kV. For quantitative measurement of distance between organelles, the closest point between *T. gondii’s* organelles and ER membrane was measured using ImageJ and was performed on 47 representative electron micrographs at the same magnification for accurate comparison between organelles

### Statistical Analysis

Statistical analyses were performed by Student’s t-test using GraphPad PRISM version 9. Error bars shown represent mean ± SD (standard deviation) of at least three independent biological replicates. Unpaired two tailed t test performed in all comparisons.

## Acknowledgments

The authors thank Dr. Muthugapatti Kandasamy and the Biomedical Microscopy Core of the University of Georgia for the use of the microscopes. The CTEGD Cytometry Shared Resource Laboratory provided access and training to state-of-the-art flow cytometry analyzers. We would like to thank David Sibley for the generous gift of the mouse SERCA antibody and the plasmid for SERCA expression. This work was supported by the U.S. National Institutes of Health grants AI128356, AI154931 and AI174600 to SNJM and R01AI166921 to IC.

