## Supplemental tables and Figures for "Calcium transfer from the ER to other organelles for optimal signaling in *Toxoplasma gondii*"

**Zhu-Hong Li et al.**

Supplemental Material

**Supplemental Table 1: Primers used in this study**

| Primer Number | Sequence | Purpose |
|---|---|---|
| 1: SERCA_5'UTR_F | ttctgcagatatccatcacactggcCCACCTCCTCGGCGGTTCTCTCTCG | Amplify 5'UTR of *TgSERCA* |
| 2: SERCA_5'UTR_R | aggtttcgtgctgCGCTGCGTCTCCGAAGATAAGCCGAAC | |
| 3: DHFR+T7S4_SERCA_F | cggagacgcagcgCAGCACGAAACCTTGCATTCAAACC | Amplify DHFR+T7S4 cassette |
| 4: DHFR+T7S4_SERCA_R | ttgacaggtccatGGTTGAAGACAGACGAAAGCAGTTG | |
| 5: SERCA_3'HR_F | tctgtcttcaaccATGGACCTGTCAAACGAGAAAGCCG | Amplify the 3' homologous region of *TgSERCA* for promoter insertion |
| 6: SERCA_3'HR_R | gggccctctagatgcatgctcgagcGGAGACTCTGAATGAGTGAACACGAG | |
| 7: SERCA-LIC-F | TACTTCCAATCCAATTTAATGCCGACGATCCCTGCTCCTT | Amplify 3' *TgSERCA* for LIC into plic-3HA |
| 8: SERCA-LIC-R | TCCTCCACTTCCAATTTTAGCCTGCAGCTTGCGCAGCTG | |
| 9: R-6f-AvrII-NoSC | CCTAGGCTTCGCTGTCATCATTTGTACA | Make SOD2-Gcamp6 construct |
| 10: F-SOD2-BglII | AGATCTATGTCCATCACAGCTGTCCTAGTGCCAG | |
| 11: R-SOD2-4-Gcamp6 | TGAGAACCCATGGCGTTTGTGGAGAAACAGTGGGC | Make SOD2-Gcamp6 construct |
| 12: F-Gcamp6-4-SOD2 | CCACAAACGCCATGGGTTCTCATCATCATCATCATC | |
| 13: XmaI_SERCA_PNP_F | 5' ACGTCCCGGGTGCCATCGTGAGAAAGCTCGCG 3' | Amplify *TgSERCA* to prepare recombinant protein for Ab production |
| 14: HindIII_SERCA_PNP_R: | 5' ACGTAAGCTTGTTGTCGTCTGCGAGAACCATG 3' | |

**Supplemental video legends:**

**Supplemental Video 1.** IFA of intracellular parasites labeled with the mitochondria marker αTom40 (green) antibody, and the ER labeled with the αTgERC antibody (red). Image acquisition using Zeiss Elyra Super resolution microscope and 3D visualizations using Imaris version 10.1.

**Supplemental Video 2.** Imaris 3D optimal visualization of extracellular parasites labeled with the αTom40 (green) antibody and the ER labeled with the αTgERC antibody (red).

**Supplemental Video 3.** ER membrane contacts sites with the plant like vacuolar compartment (PLVAC). Imaris 3D visualization of immunofluorescence of extracellular tachyzoites. PLVAC was labeled with αVP1 antibody (green) and ER was labeled with αTgERC antibody (red)

**Supplemental Figures**

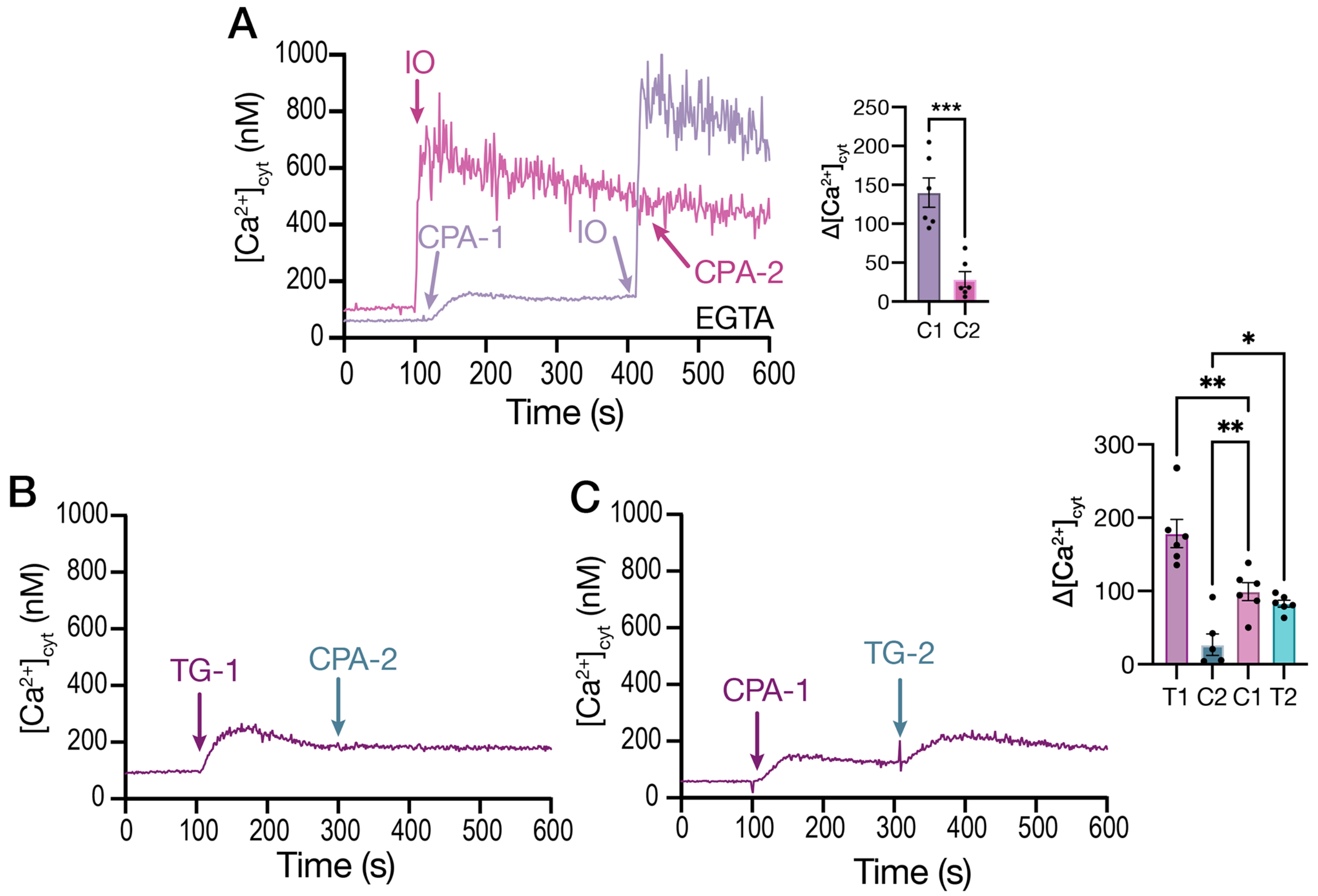


**Figure S1: Intracellular Calcium pools.** *T. gondii* tachyzoites loaded with Fura-2 were used for these measurements. The suspension was in Ringer buffer with 100 µM EGTA. **A**, Cyclopiazonic acid (CPA), 10 µM was added at 100 seconds (CPA-1) followed by Ionomycin 1 µM at 400 sec (*purple trace*). CPA-2, *pink trace*: 1 µM IO was added at 100 sec followed by CPA 10 µM at 400 sec. The bar graph shows the statistical analysis of cytosolic $\Delta[Ca^{2+}]$ measured after the addition of CPA first (C1), compared with the response to CPA after addition of IO (C2), based on data from more than three independent biological experiments. **B**, TG 1 µM was added at 100 sec followed by CPA at 300 sec. **C**, CPA 10 µM was added at 100 sec followed by TG 1 µM at 300 sec. The bar graph shows the analysis of the cytosolic $\Delta[Ca^{2+}]$ after the addition of TG (T1 and T2) or CPA (C1 and C2) from more than three biological experiments. Data are presented as mean ± SD. *p value*: unpaired two tailed t test performed in all comparisons. *, $p \leqslant 0.05$. **, $p \leqslant 0.01$. ***, $p \leqslant 0.001$. ****, $p \leqslant 0.0001$.

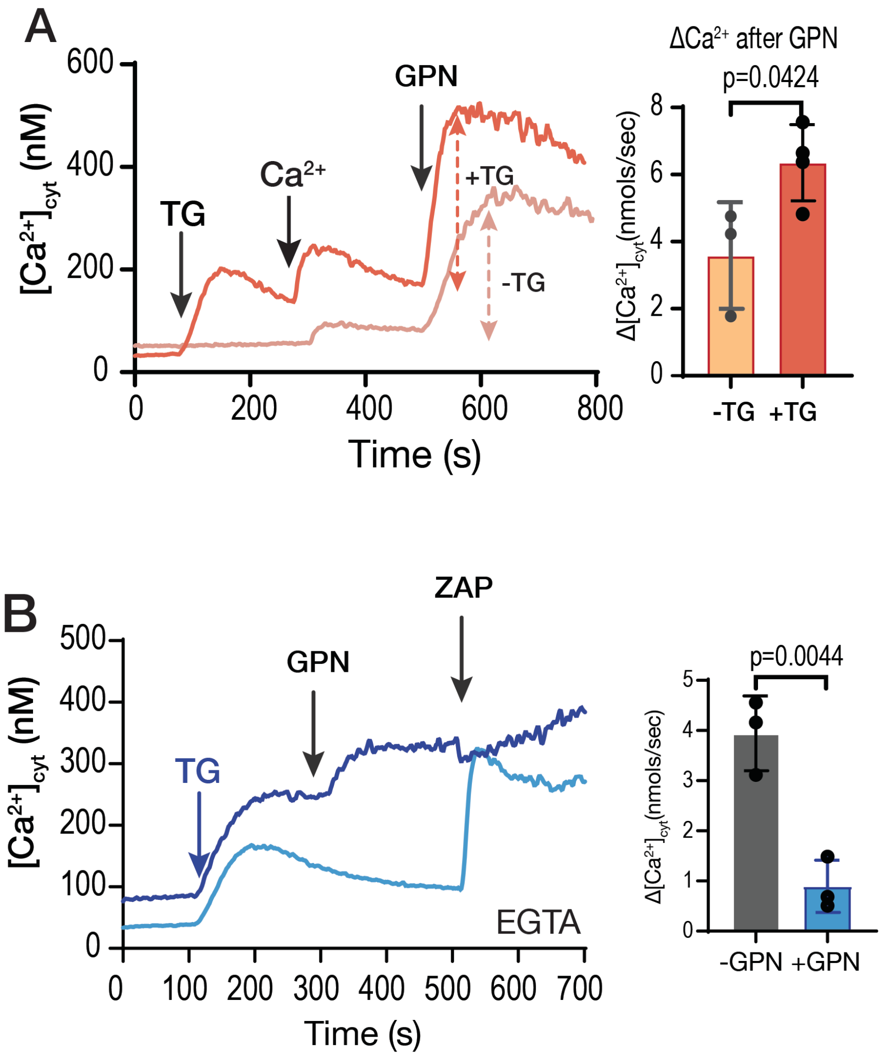


**Figure S2: Intracellular Calcium pools**. *T. gondii* tachyzoites loaded with Fura-2 were used for these measurements. The suspension was in Ringer buffer with 100 μM EGTA. **A**, 1 μM Thapsigargin (TG), was added at 100 seconds followed by 1.8 mM $Ca^{2+}$ at 300 sec and 40 μM GPN at 500 sec (*orange trace*). The *peach trace* shows the same additions of $Ca^{2+}$ and GPN but no TG addition. The bar graph shows the statistical analysis of the rate of $\Delta[Ca^{2+}]_{cyt}$ per second obtained after the addition of GPN. **B**, Similar experimental set-up: 1 μM TG was added at 100 sec, followed by 40 μM GPN (*dark blue trace*) and 100 μM Zaprinast at 500 sec. The bar graph shows the analysis of the rate of the cytosolic $\Delta[Ca^{2+}]$ change after the addition of Zaprinast from more than three biological experiments. Data are presented as mean ± SD. *p value*: unpaired two tailed t test performed in all comparisons.

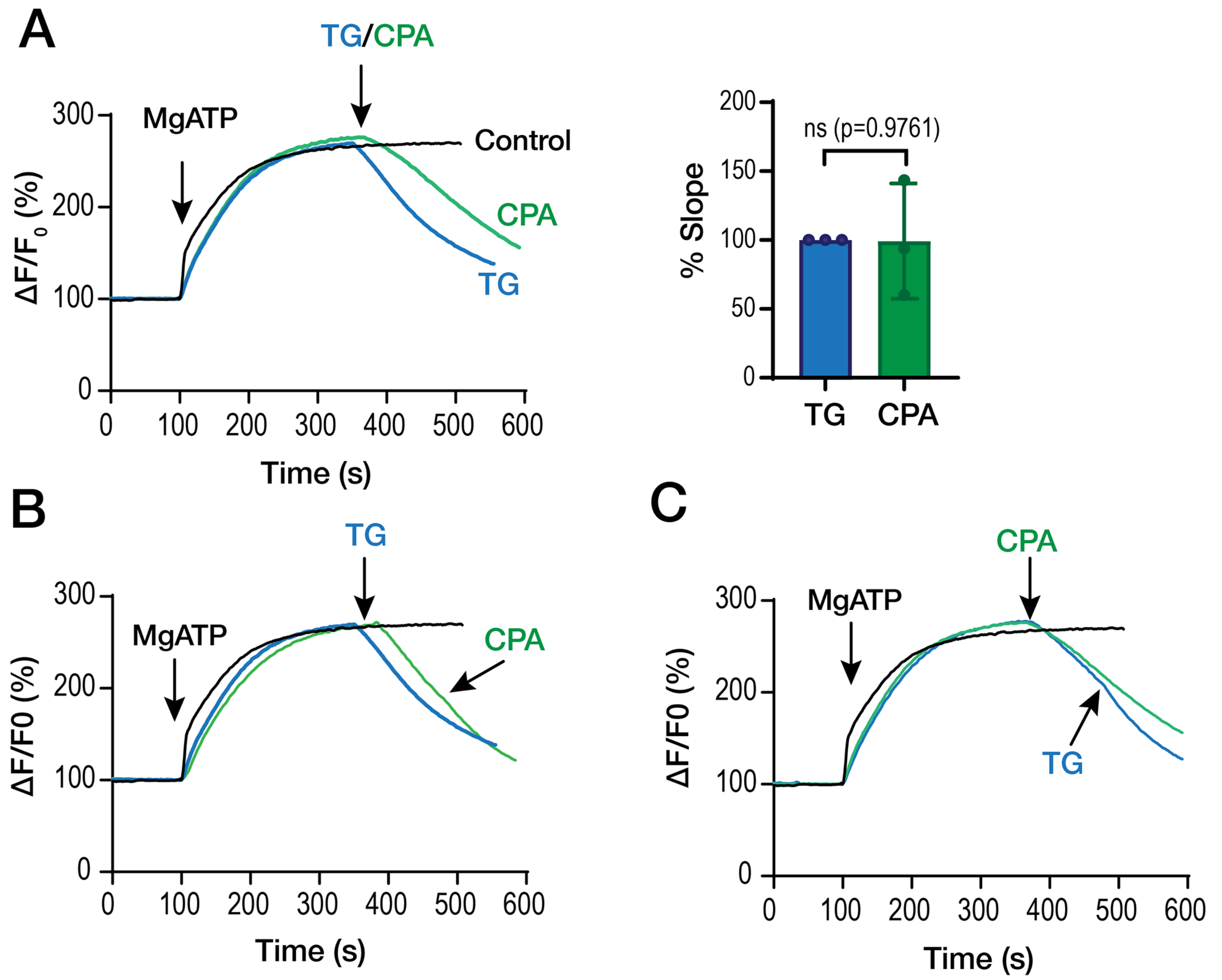


**Figure S3: Inhibition of SERCA activity by thapsigargin (TG) and cyclopiazonic acid (CPA).** *T. gondii* tachyzoites loaded with Mag-Fluo4-AM and permeabilized with digitonin were used for these measurements. **A**, 125 µM MgATP was added at 100 seconds, followed by the addition of DMSO (control) or 1 µM TG or 10 µM CPA at 350 seconds. The slope of the fluorescence decrease after CPA addition was normalized to the slope of the fluorescence decrease after TG addition, which was set to 100% in the bar graph. **B,** MgATP was added at 100 sec followed by TG at 350 seconds for both runs, CPA at 500 sec was added to the experiment represented by the green trace. **C,** MgATP was added at 100 sec followed by CPA at 350 seconds for both runs, TG was added at 500 Seconds to the experiment represented by the blue trace.

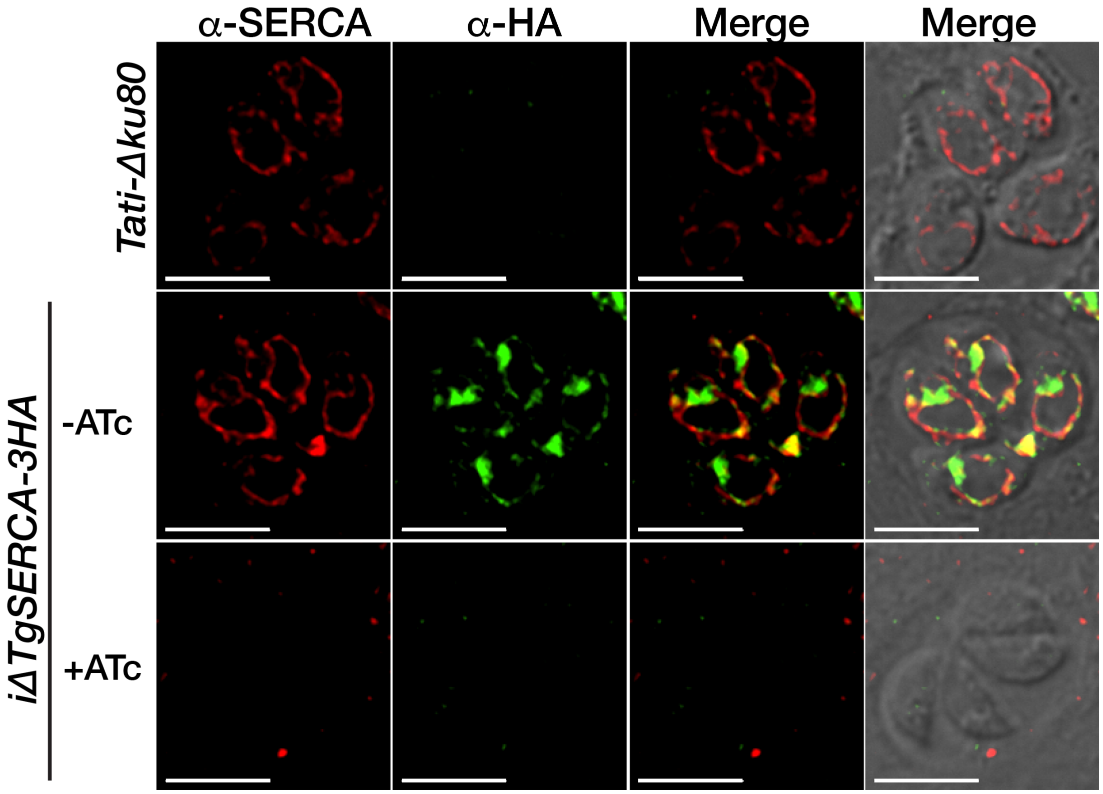


**Figure S4: Regulation of the expression of TgSERCA**. IFAs of *TatiΔku80* or *iΔTgSERCA-3HA* (±*ATc*). The mouse monoclonal antibody αHA was used at 1:25 dilution (green signal). The Guinea pig antibody against TgSERCA was used at 1:500 dilution (red signal). The HA signal partially co-localizes with the TgSERCA signal. Both signals disappear in the *iΔTgSERCA-3HA* mutant when cultured with ATc for 24 hours.

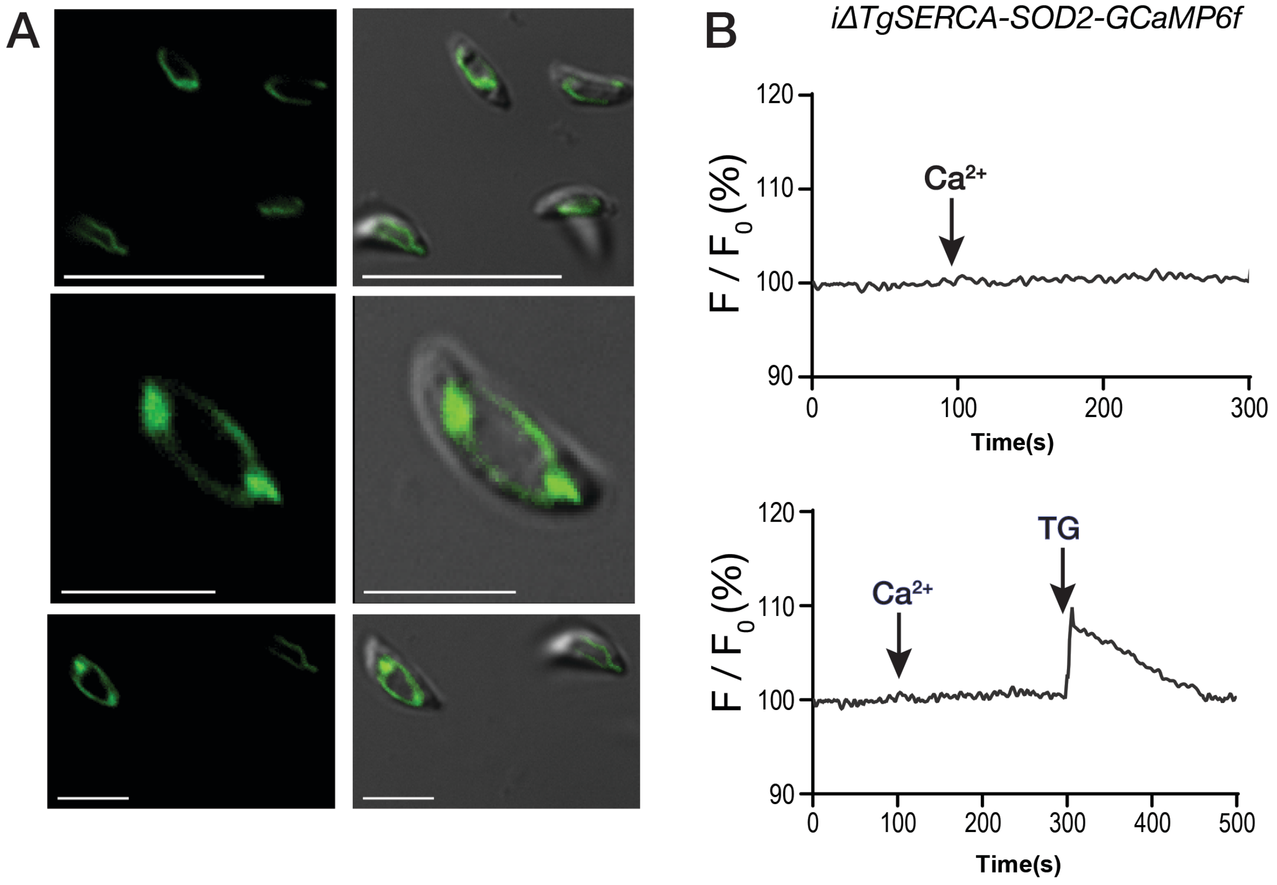


**Figure S5:** Mitochondrial localization of the GCaMP6f. **A,** *T. gondii* tachyzoites of the *iΔTgSERCA*-SOD2-GCaMP6f clonal mutant live show localization of the fluorescence signal in the mitochondrion. **B**, Live intact *iΔTgSERCA*-SOD2-GCaMP6f mutant parasites in suspension show the lack of response to the addition of extracellular $Ca^{2+}$. The lower trace shows a similar experiment with the addition of TG at 300 sec. Parasites were in suspension in Ringer buffer.

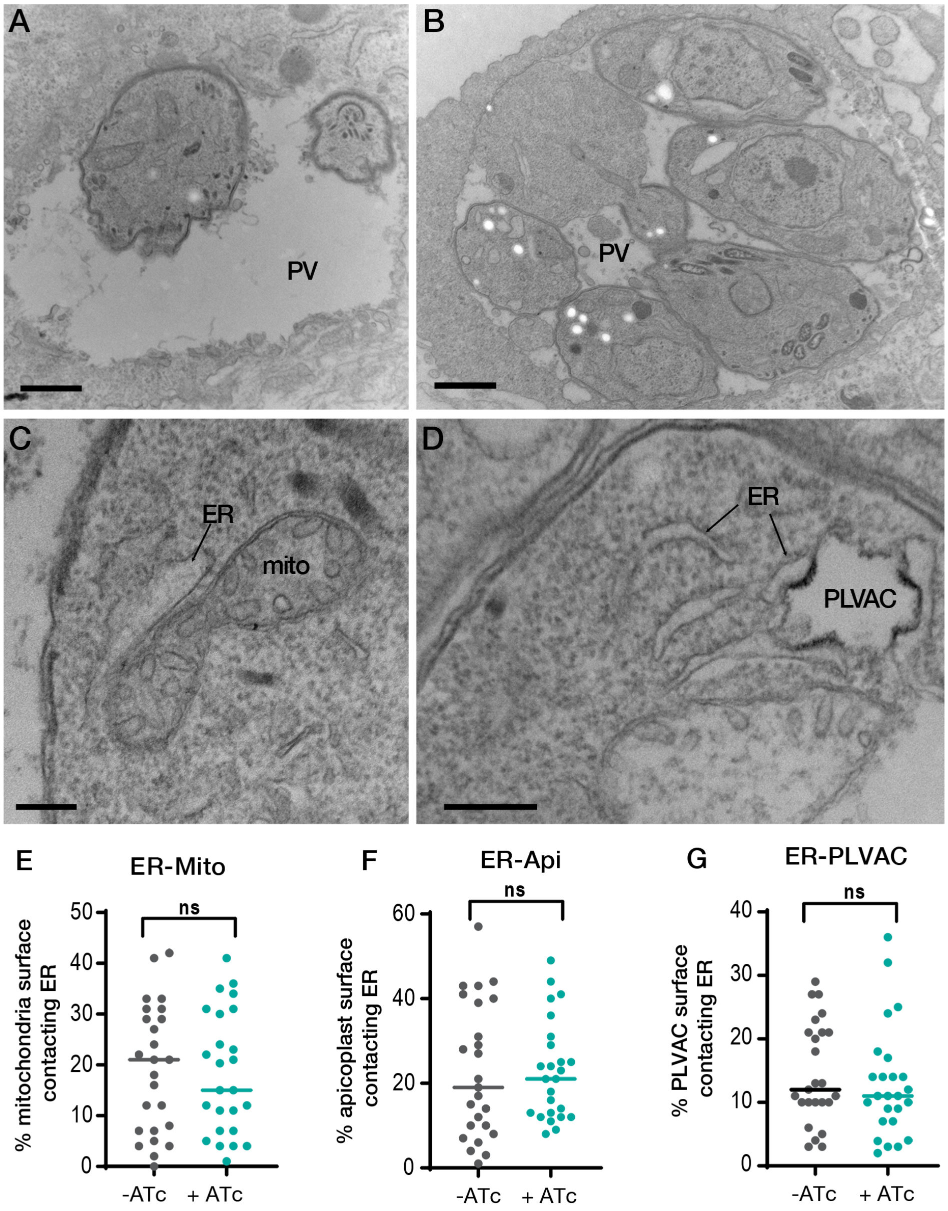


**Figure S6:** Electron microscope images of the *iΔTgSERCA* mutant treated with ATc for 24h. **A**, Representative images of the *iΔTgSERCA* mutant highlighting a large empty PV. Size bar is 500 nm. **B**, Representative image of the *iΔTgSERCA* mutant with a large residual body inside the PV. Bar represents 500 nm. **C**, The *iΔTgSERCA* mutant treated with ATc still showed contacts between the ER and mitochondria. Bar represents 100 nm. **D**, The *iΔTgSERCA* mutant treated with ATc still showed contacts between the ER and the PLVAC. Size bar is 100 nm. **E-G,** quantitative assessment of the contact area between ER and organelles (mitochondrion, apicoplast, or PLVAC), length of the limiting membrane of the organelle in contact with ER tubules at a distance less than 30 nm. This was measured and divided by the total length of the limiting membrane of the organelle. A total of 47 to 85 sections was analyzed for each population of organelles. **E**, Comparison of contacts measurements for *iΔTgSERCA* ± ATc for ER-mitochondria. **F**, Comparison of contacts measurements for *iΔTgSERCA* ± ATc for ER-apicoplast, **G**, Comparison of contacts measurements for *iΔTgSERCA* ± ATc for ER-PLVAC. All *p* values were calculated by two-tail t test comparing *iΔTgSERCA* ± ATc. *p* values ER-mito: 0.403; ER-Api: 0.492; ER-PLVAC: 0.244
